# HOS15-mediated turnover of PRR7 enhances freezing tolerance

**DOI:** 10.1101/2024.06.20.599783

**Authors:** Yeon Jeong Kim, Woe Yeon Kim, David E. Somers

## Abstract

Arabidopsis *PSEUDO RESPONSE REGULATOR7* (*PRR7*) is a core component of the circadian oscillator which also plays a crucial role in freezing tolerance. PRR7 undergoes proteasome-dependent degradation to discretely phase maximal expression in early evening. While its transcriptional repressive activity on downstream genes is integral to cold regulation, the mechanism of the conditional regulation of the PRR7 protein activity is unknown. We used double mutant analysis, protein interaction and ubiquitylation assays to establish that the ubiquitin ligase adaptor, *HIGH EXPRESSION OF OSMOTICALLY RESPONSIVE GENE 15* (*HOS15*), controls the protein accumulation pattern of PRR7 through direct protein-protein interactions. Freezing tolerance and electrolyte leakage assays show that PRR7 enhances cold temperature sensitivity, supported by ChIP-qPCR at *C-REPEAT BINDING FACTOR* (*CBF*) and *COLD REGULATED 15A* (*COR15A*) promoters where PRR7 levels were higher in hos15 mutants. We establish that HOS15 mediates PRR7 protein turnover through enhanced ubiquitylation at low temperature in the dark. Under the same conditions, increased PRR7 association with the promoter regions of *CBFs* and *COR15A* in *hos15* correlates with decreased *CBF1* and *COR15A* transcription and enhanced freezing sensitivity. We propose a novel mechanism whereby HOS15-mediated regulation of PRR7 provides an intersection between the circadian system and other cold acclimation pathways leading to freezing tolerance through upregulation of *CBF1* and *COR15A*.

## Introduction

Plants have evolved an internal timekeeping mechanism comprised of molecular processes in response to the regular daily oscillations of environmental cues (Linde *et al*., 2017; Michael, 2022; Petersen *et al*., 2022). This internal circadian oscillator coordinates a self-sustaining ∼24-hour sinusoidal rhythm of gene expression and cellular metabolism, allowing the plant to respond appropriately to fluctuations in ambient light, temperature, humidity, and nutrition (Inoue *et al*., 2017; Xu *et al*., 2022). Diurnal cycles of light and temperature entrain the circadian system, which adjusts the timing and strength of output responses to modulate plant physiology and development (Oravec & Greenham, 2022; Wang *et al*., 2022; Xu *et al*., 2022; Xu *et al*., 2023). During the transmission of input signals to output pathways, the circadian oscillator regulates the expression of output genes through transcriptional control, and *PSEUDO-RESPONSE REGULATOR* (*PRR*) genes play essential roles (Farré & Liu, 2013).

*PRR* genes are a family of transcriptional repressors consisting of five members, the morning-phased genes *PRR9* and *PRR7* and the evening-phased genes *PRR5*, *PRR3*, and *TOC1(PRR1)* in Arabidopsis (Farré & Liu, 2013). The transcript levels of *PRR9/7/5/3/1* peak sequentially at different times during the day and night, each with a narrow peak expression window, which is also mirrored in their protein expression patterns (Fujiwara *et al*., 2008; Nakamichi, N. *et al*., 2010). *PRR7* is an important component of the central oscillator as well as playing a central role in transmitting phase-specific control to key abiotic genes (Farré *et al*., 2005; Nakamichi *et al*., 2005; Salomé & McClung, 2005; Nakamichi *et al*., 2009; Salomé *et al*., 2010; James *et al*., 2012; Liu *et al*., 2013; Kolmos *et al*., 2014; Liu *et al*., 2015; Yuan *et al*., 2021).

A role for *PRR7* in cold tolerance is seen in the increased freezing stress of *prr9prr7* and *prr9prr7prr5* mutants (Nakamichi *et al*., 2009; Wang *et al*., 2017). Additionally, PRR7 associates with the promoter of the cold tolerance genes *C-REPEAT/DRE BINDING FACTOR1* (*CBF1*), *CBF2*, and *CBF3* (Nakamichi *et al*., 2012; Liu *et al*., 2013; Liu *et al*., 2015), and the *prr9prr7prr5* triple mutant shows high expression level of *CBFs*, consistent with strong freezing tolerance (Nakamichi *et al*., 2009).

The regulation of *PRR7* by the environmental input signals involves transcriptional (Frank *et al*., 2018; Nakamichi, 2020), post-transcriptional (Wang *et al*., 2022), and post-translational mechanisms (Farré & Kay, 2007; Fujiwara *et al*., 2008). Light induces the accumulation of *PRR7* mRNA in the morning by relieving repression by other clock components, whereas darkness activates the transcriptional repressors of *PRR7* (Nakamichi, 2020) and promotes PRR7 degradation via a proteasome-dependent pathway (Farré & Kay, 2007). The rapid decline in PRR7 protein abundance in the dark is an important mechanism that shapes a robust oscillation of the protein and restricts the activity of PRR7 to late in the day/early evening. While the F-box protein ZEITLUPE (ZTL) and related proteins (LKP2, FKF1), target TOC1 and PRR5 for degradation (Más *et al*., 2003; Kiba *et al*., 2007; Baudry *et al*., 2010; Wang *et al*., 2010), this family is unlikely to regulate PRR7 protein (Fujiwara *et al*., 2008), and factors responsible for PRR7 turnover remain unknown.

HOS15 is a DWD (DDB1-binding WD40) motif-containing protein (Lee *et al*., 2008) that recruits HISTONE DEACETYLASE 2C (HD2C) to the DDB1-cullin4 machinery and mediates the ubiquitination and degradation of HD2C at 4°C (Park *et al*., 2018). At low temperatures, HOS15 also promotes the degradation of GIGANTEA (GI), a multifunctional protein which regulates transcription, photoperiodic flowering and stabilizes ZTL in the circadian clock (Kim *et al*., 2007; Nohales *et al*., 2019; Ahn *et al*., 2023). Accumulating evidence also suggests that *HOS15* is a positive regulator of freezing tolerance and that the underlying mechanisms are based on functions involved in histone deacetylation and ubiquitination at low temperatures (Zhu *et al*., 2008; Park *et al*., 2018).

In this study, we tested the relationship between HOS15 and PRR7 in the regulation of the cold response pathway in Arabidopsis. We found that HOS15 interacts with PRR7 and facilitates its ubiquitination at low temperatures. The HOS15-mediated PRR7 turnover operates at low temperatures in the dark, resulting in an acute decrease in PRR7 protein abundance at dusk and subsequent derepression of two independent targets of the *ICE-CBF-COR* cold tolerance pathway (Hwarari *et al*., 2022), *CBF1* and *COR15A,* to enhance freezing tolerance. Our results demonstrate a novel role of HOS15 to refine the waveform of PRR7 protein abundance during entrainment under moderately low temperatures to regulate cold acclimation.

## Material and methods

### Plant materials, growth condition, and chemical treatment

*PRR7pro::FLAG-PRR7-GFP/*Col-0 (*F7G*) and *PRR7pro::FLAG-PRR7-GFP/hos15-2* (*F7Ghos15-2*) seedlings were generated by crossing *F7G* (Nakamichi, Norihito *et al*., 2010) with *hos15-2* (GK_785B10). The *hos15-2prr7-3* (*prr7-3*, SALK_030430) and *prr9-1prr7-3* (*prr9-1*, SALK_007551) double mutants were obtained by crossing single mutants. The *35S::TAP-PRR7* transgenic line was constructed by transforming pN-TAPa-PRR7 into Col-0, and *35S::TAP-PRR7/*Col-0 and *35S::TAP-PRR7/hos15-2* lines were selected among F_3_ progeny of cross of *35S::TAP-PRR7* T_1_ plant and *hos15-2*.

For Arabidopsis growth, seeds were sterilized and kept in the dark at 4°C for 4 d before sowing. Seedlings were grown under a 12 h/12 h light/dark cycle (12L/12D; white fluorescent light, 50 μs^-1^) on MS medium (Caisson Laboratories Inc) supplemented with 3% sucrose and 0.8% agar for the period indicated in each figure legend. For low temperature acclimation, the seedlings on MS plates were transferred to 15°C and grown under the same light regime for 3 d. Subsequent cycloheximide treatment was performed by transferring seedlings to MS liquid medium containing 100 μ 0.01% Triton X-100 and incubating for 10 h with shaking. For MG132 treatment, the seedlings were M MG132 for 5 x 1 μm and shaken for 13 h. For freezing tolerance and ion leakage assays, the seeds were sown on soil, stratified for 4 d, and grown in 12L/12D (100 μmol m^-2^ s^-1^) at 22°C for 24 d and at 15°C for an additional 7 d.

### Plasmid construction

DNA regions corresponding to HOS15 full length (1-613 aa), N-terminal region (1-248 aa), and C-terminal region (249-613 aa) were amplified from cDNA with or without STOP codon and cloned into pENTR2B (Invitrogen) or pENTR/D-TOPO (Invitrogen). *HOS15* expression clones were combined with entry clones and pCsVMV-HA-C1300, pDEST32, or pSITE-BiFC-nEYFP-N1 using LR clonase (Invitrogen). Full-length PRR7 cDNA was amplified from the previously generated expression clone *pCsVMV-GFP-C1300-PRR7* (Wang *et al*., 2013) and cloned into pENTR2B, followed by subcloning into pDEST22 and pSITE-BiFC-cEYFP-C1 vectors. For *35S::TAP-PRR7* construction, pENTR2B-PRR7 was created and combined with pN-TAPa vector (Rubio *et al*., 2005) by LR recombination. The primers used for plasmid construction are described in Table S2.

### Coimmunoprecipitation

For transient *N. benthamiana* expression, four-week-old *N. benthamiana* leaves were infiltrated with Agrobacteria GV3101 strain containing pCsVMV-HA-C1300-*HOS15* and/or pCsVMV-GFP-C1300-*PRR7* and collected after 3 d. *N. benthamiana* leaves and Arabidopsis seedlings were ground using liquid nitrogen and total proteins were extracted using cold IP buffer (50 mM Tris–Cl, pH 7.5, 150 mM NaCl, 0.5% Nonidet P-40, 1 mM EDTA, 2 mM dithiothreitol, 2 mM sodium fluoride, 2 mM sodium vanadate, 1 g/ml pepstatin, 5 μM ALLN) from 1.5 μml of tissue powder. Protein G-agarose resin pretreated with mouse monoclonal anti-GFP (Thermo Scientific, A11120) overnight was incubated with clarified supernatant for 2 h at 4°C. After centrifugation, immune complexes were rinsed with cold IP buffer at least six times at 4°C and released from beads by heating at 80°C for 2 m. The eluted proteins were separated on 8% SDS-PAGE gel (37.5:1 acrylamide:bis-acrylamide) and probed with anti-HA (Roche, 3F10), anti-GFP (Abcam, ab6556) and anti-HOS15 (Lee *et al*., 2012) antibodies.

### Yeast two-hybrid

GAL4 DNA-binding domain-tagged HOS15 and GAL4 activation domain-tagged PRR7 were co-transformed into AH109 containing *HIS3* and *ADE2* selectable genes controlled by GAL4-binding sites using the lithium chloride method (Elble, 1992). Transformed cells were selected on a SD medium (Difco) deficient in leucine and tryptophan (SD-Leu-Trp), and individual colonies were subcultured on a SD medium lacking histidine, leucine, and tryptophan for 7 d. For the β-galactosidase assay, three individual colonies were cultured in SD-Leu-Trp and incubated with o-nitrophenyl-β-D-galactopyranoside (Sigma). The resulting absorbance values at 420 nm were normalized to protein concentration quantified by Bradford assays, and arbitrary β-galactosidase units were determined as follows: absorbance value at 420 nm × 1.7)/(0.0045 × incubation time × volume of extract × protein concentration in mg/ml.

### *In vivo* ubiquitination

Corresponding Arabidopsis seedlings were vacuum treated with DMSO 139 or MG132, and total proteins were extracted from 1 ml of tissue powder using extraction buffer (50 mM Tris–Cl, pH 7.5, 150 mM NaCl, 0.5% Nonidet P-40, 1 mM EDTA, 2 mM dithiothreitol, 2 mM sodium fluoride, 2 mM sodium vanadate, 1 mM phenylmethylsulfonyl fluoride, 5 μg/ml leupeptin, 1 μg/ml aprotinin, 1 μg/ml pepstatin, 5μg/ml antipain, 5 μg/ml chymostatin, 50 μM MG132, 50 μM MG115, and 50 μM ALLN, 10 nM Ub aldehyde, and 10 mM N-ethylmaleimide). Human IgG-Sepharose beads (Cytiva) were added to the protein extracts, and after 2 h incubation, the beads were washed three times with wash buffer (50 mM Tris–Cl, pH 7.5, 150 mM NaCl, 0.5% Nonidet P-40, 1 mM EDTA, 2 mM dithiothreitol, 2 mM sodium fluoride, 2 mM sodium vanadate, 10 nM Ub aldehyde, and 10 mM N-ethylmaleimide) and once with wash buffer without 10 nM Ub-aldehyde, and 10 mM N-ethylmaleimide. The immune complexes were released from the resin by HRV-3C protease (Pierce, Cat. No. 88946) treatment at 4°C for 2 h and loaded onto 8% SDS-PAGE gel. The membranes were probed with anti-myc (Sigma, M4439) and anti-Ub (Santa-Cruz, sc-8017).

### Freezing tolerance and ion leakage assays

For freezing tolerance evaluation Col-0, *hos15-2*, *prr7-*3, *hos15-2prr7-3*, and *prr9-1prr7-3* were cultivated on soil in 12L/12D at 22°C for 24 d followed by 12L/12D at 15°C for one week. These plants were then sequentially exposed at ZT9 to 4°C for 1 h, then harvested either after −4°C or −6°C treatments after being sequentially exposed to 0°C for 1 h, −2°C for 1h, and then either −4°C only for 1h, or then −6°C for 1 h followed by 4°C for 16 h in darkness to thaw soil. Survival rate (100 – yellow plants/green plants) was measured with at least 19 plants in each trial 7 d after growth recovery in 12L/12D at 15°C. For the ion leakage assay, plants were grown under the same condition as for the freezing tolerance assay, except an additional −8°C treatment for 1 h. Leaf discs (approximately 1.5 cm) were collected from 10 rosette leaves of the plants at 4°C after each −2, −4, −6, and −8°C treatment, and incubated overnight in 20 ml deionized water with shaking. The conductivity in the solution was measured before and after 1 h of boiling using a Cond 330i meter and a TetraCon 325 probe (WTW, Germany). The percentage of electrolyte leakage was calculated as follows: initial conductivity / maximum conductivity x 100.

### Bimolecular fluorescence complementation assay

BiFC constructs were transformed into Agrobacteria GV3101 and infiltrated into four-week-old *N. benthamiana* leaves. Samples were excited with 514-nm and 543-nm lasers for YFP-fused proteins and H2B-RFP, respectively, and fluorescence was collected at 522-555 and 585–615 emission filters. Images were captured using a Nikon A1+ confocal microscope with a 100× water immersion objective (NA = 1.4), using the 3× confocal zoom. Images were processed using Nikon NIS-Elements software and ImageJ v1.53.

### RNA extraction and Quantification Real-Time PCR

Total RNA was extracted with Trizol reagent (Invitrogen) and treated with TURBO DNA-free kit (Invitrogen) according to the manufacturer’s protocol. First-strand cDNA was synthesized with 1 μ total RNA using SuperScript^TM^ IV reverse transcriptase (Thermo Fisher Scientific), and the fluorescence curve were determined using PowerUp™ SYBR™ Green Master Mix (Thermo Scientific) and QuantStudio^TM^ 5 Real-Time PCR instrument (Applied Biosystems). Relative transcript abundance was calculated by normalization to that of *ACTIN* (AT3G18780). Thermocycling conditions were as follows: 94◦ C for 2 min; followed by 44 cycles of 94◦ C for 15 s and 55◦ C for 34 s. The information on the primers used for ChIP-qPCR is provided in Table S2 (Jiang *et al*., 2017).

### Chromatin immunoprecipitation

ChIP-qPCR was performed according to the following protocol (Yamaguchi *et al*., 2014) with minor modifications (Liu *et al*., 2013). Briefly, the collected seedlings in 50 ml tubes covered with aluminum foil were cross-linked by vacuum infiltration in 1X PBS solution containing 1% formaldehyde for 8 x 1 minutes on ice, followed by 1.5 m vacuum in cold 0.125 M glycine. After three washes with cold 1X PBS buffer, the tissues were collected and finely ground. For nuclei isolation, 1 ml of tissue was homogenized with 1X Nuclei PURE Lysis Buffer (NIBA buffer supplemented with 2 mM dithiothreitol, 2 mM sodium fluoride, 2 mM sodium vanadate, 1 mM phenylmethylsulfonyl fluoride, 5 μg/ml leupeptin, 1 μg/ml aprotinin, 1 μg/ml pepstatin, 5 μg/ml antipain, 5 μg/ml chymostatin, 50 μM MG132, 50 μM MG115, and 50 μM ALLN; Nuclei PURE prep nuclei isolation kit, Sigma), and clarified by filtration through Miracloth (Millipore). The nuclei pellet was collected by centrifugation and resuspensed in 1X NIBA buffer containing 0.3 % Triton-X100, and incubated in nuclei lysis buffer (50 mM Tris-HCl pH 8.0, 10 mM EDTA pH 8.0, 1 % SDS). ChIP dilution buffer (16.7 mM Tris-HCl pH 8.0, 167 mM NaCl, 1.2 mM EDTA, 0.01 % SDS) was added to the nuclei lysate and lysed by two 30 s-sonication pulses using a Covaris S220 Focused-ultrasonicator (peak incident power 140 watts, duty factor 10%, cycles per burst 200). After centrifugation, the clarified supernatant was mixed with Dynabeads Protein G (Invitrogen), which was pre-incubated with anti-GFP (Abcam, ab6556) overnight, at 4°C for 2 h and washed five times as described in previous study (Liu *et al*., 2013). Reverse cross-linking was performed by incubating the beads repeatedly at 65°C for 30 min, and recovered DNA was purified using the QIAquick PCR Purification kit (Qiagen). The relative abundance of DNA in the resulting elute was quantified by normalization to the amount in the input DNA.

## Results

### *HOS15* reduces PRR7 abundance at low temperature

HOS15 regulates the stability of the clock protein GI at 16°C (Ahn *et al*., 2023) and interacts with the Evening Complex (Park *et al*., 2019). Both HOS15 and PRR7 are involved in the cold response pathway (Zhu *et al*., 2008; Nakamichi *et al*., 2012; Liu *et al*., 2013; Liu *et al*., 2015; Park *et al*., 2018; Ahn *et al*., 2023), so we tested the effect of a HOS15 deficiency on PRR7 protein abundance at both 22°C and 15°C (Fig. 1a,b). Under these conditions, PRR7 protein abundance peaks around dusk at 22°C (Fujiwara *et al*., 2008; Nakamichi, N. *et al*., 2010), with no difference between wild type (WT) and mutant, while at 15°C levels are significantly higher during the night in *hos15-2* at ZT13 (zeitgeber time) and ZT21. *PRR7* transcript levels are comparable in the two backgrounds at 15°C (Fig. 1c), indicating that the higher PRR7 levels in the mutant occurs post-transcriptionally (Fig. 1c). When plants were grown in light/dark warm (22°C)/cold (15°C) conditions similar increased PRR7 levels in the *hos15* mutant in the cold and dark were observed (Fig. S1).

**Figure 1.**
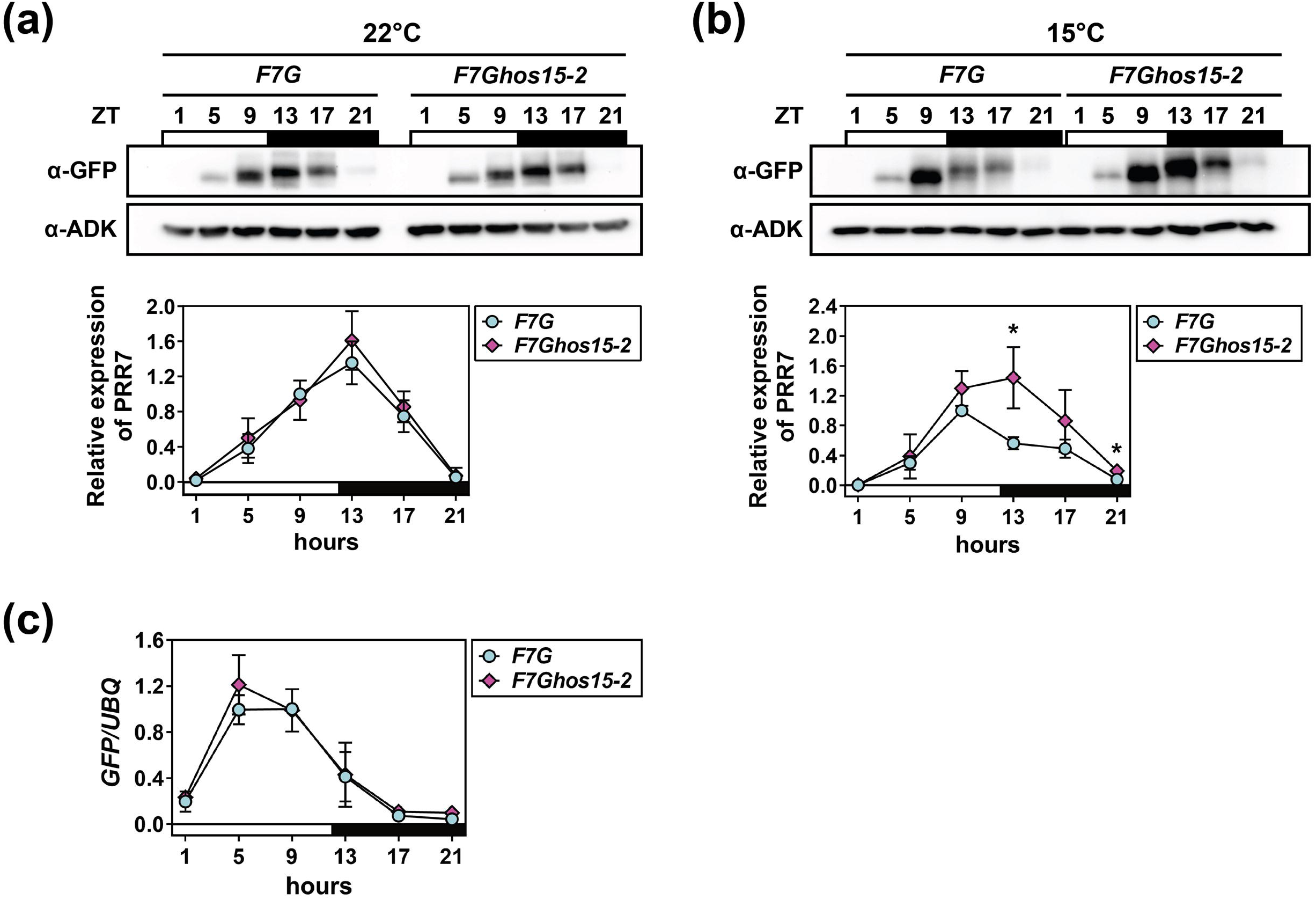
PRR7 abundance in the dark at low temperature is reduced by HOS15. (a) PRR7 protein abundance at 22°C. *PRR7pro::FLAG-PRR7-GFP* (*F7G*) and *PRR7pro::FLAG-PRR7-GFP/hos15-2* (*F7Ghos15-2*) seedlings were grown in a 12 h light/12 h dark (12L/12D) cycle at 22°C for 11 d and harvested at the times (h) indicated. PRR7-GFP protein levels were normalized to ADK (n=3). White- and black-filled rectangles indicate light and dark time periods, respectively. Statistical analysis was performed using a two-tailed Welch’s t-test for each time point. Non-significant differences (p > 0.05) are not indicated. (b) PRR7 protein abundance at 15°C. *F7G* and F7Ghos15-2 seedlings grown in 12L/12D at 22°C for 7 d were transferred to 15°C under the same light regime and collected at the indicated ZT time points 3 d after transfer. Detection and quantitation as in (A). Asterisks indicates a difference with a p-value less than 0.05, based on a two-tailed Welch’s t-test. (c) Transcript level of the *PRR7* transgene at 15°C. The transcript levels of *F7G* seedlings used in (B) were analyzed by RT-qPCR and normalized to a *U-box* gene (*At5g15400*). Error bars indicate the standard deviation of three biological replicates. No significant statistical differences (p > 0.05) between the two samples were observed based on a two-tailed Welch’s t-test.

### *HOS15* facilitates PRR7 ubiquitination and turnover in the dark at low temperature

To assess a possible role for HOS15 in the control of PRR7 protein stability at low temperatures, we performed cycloheximide chase experiments with *F7G* and *F7Ghos15-2* plants at 22°C and 15°C (Schneider-Poetsch *et al*., 2010) beginning at peak *PRR7* transcript levels (ZT9). Subsequent incubation in either continuous light or dark at both temperatures tested the effect of light and temperature on PRR7 degradation rate. At 22°C, PRR7 turnover in both WT and mutant are very similar in both light and dark (Fig. 2a,B), while at 15°C PRR7 levels decline more slowly in the mutant only in the dark (Fig. 2c,d). These findings suggest that HOS15 facilitates PRR7 turnover specifically under cool and dark conditions.

**Figure 2.**
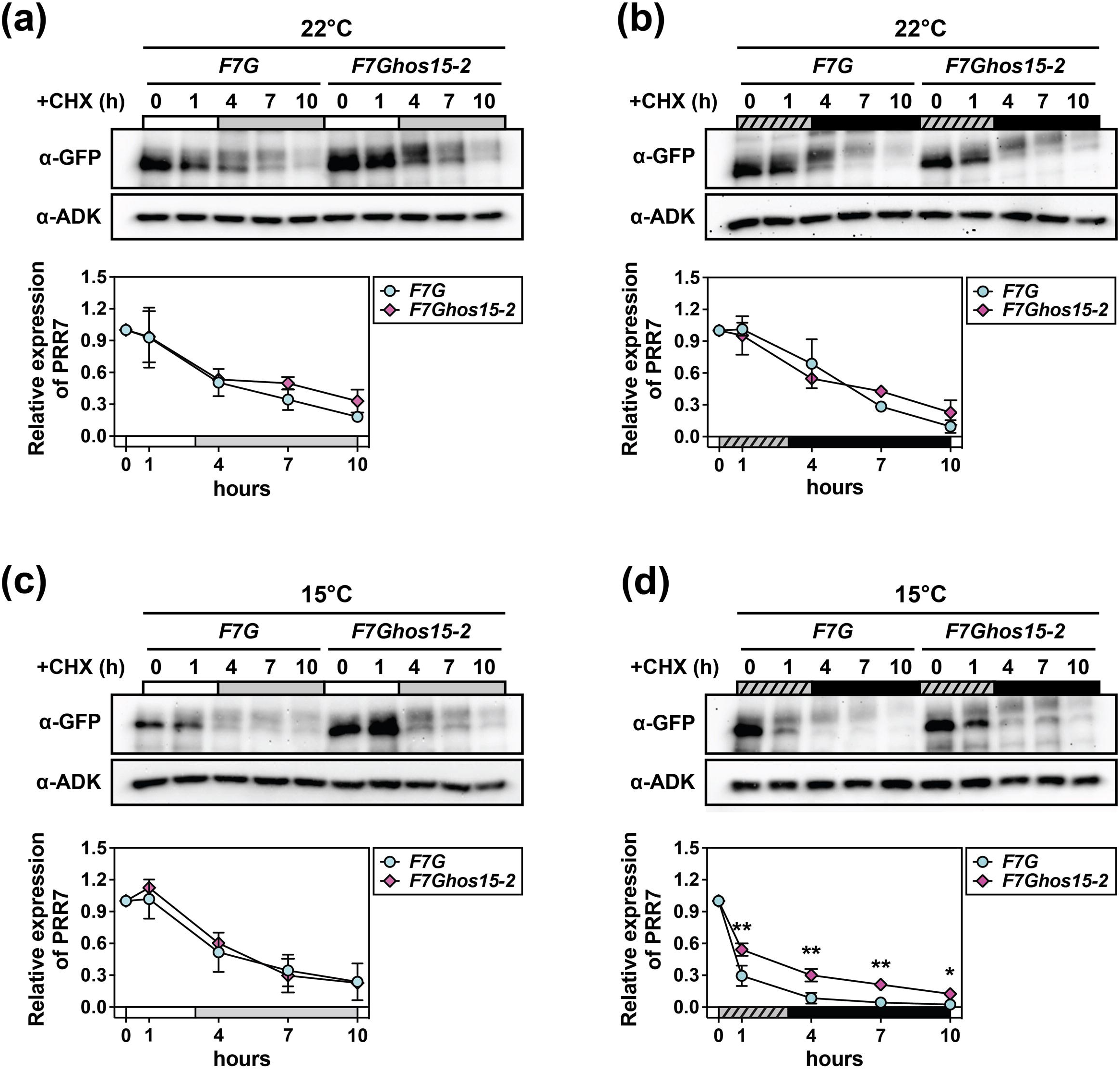
*HOS15* facilitates PRR7 turnover in the dark at low temperature. (a-d) PRR7 abundance in *F7G* and *F7Ghos15-2* seedlings after cycloheximide treatment under constant light at 22°C (a), constant dark at 22°C (b), constant light at 15°C (c), and constant dark at 15°C (d). Seedlings grown in 12L/12D at 22°C for 10 d, followed by 3 d at 22°C or 15°C, were transferred to liquid MS media containing 100 μM cycloheximide at 9 h after dawn. Seedlings were harvested at 0, 1, 4, 7, and 10 h after treatment under the indicated light and temperature conditions. Immunodetection as in Fig. 1. White, black, gray, and hatched bars indicate light, dark, subjective night, and subjective day, respectively. Mean values were scaled to the value at 0 h and shown with error bars representing the standard deviation of two (a-c) or four (d) independent trials. Asterisks indicate significant differences in a two-tailed Welch’s t-test (*, p < 0.05; **, p < 0.01). The nonsignificant differences (p > 0.05) were not depicted.

To determine whether this result derives from a direct interaction between PRR7 and HOS15, we transiently expressed HOS15 and PRR7 proteins in *N. benthamiana* leaves and performed co-immunoprecipitation assays (co-IP) (Fig. 3a). Under these conditions a substantial interaction was observed (Fig. 3a). This was further investigated at native expression levels (*F7G)* in Arabidopsis under both 15°C and 22°C and in the light (ZT9) and dark (ZT13). HOS15 preferentially co-precipitates with PRR7 at 15°C and in the dark (Fig. 3b; Fig.S2), consistent with the conditions under which PRR7 levels are reduced in the presence of HOS15.

**Figure 3.**
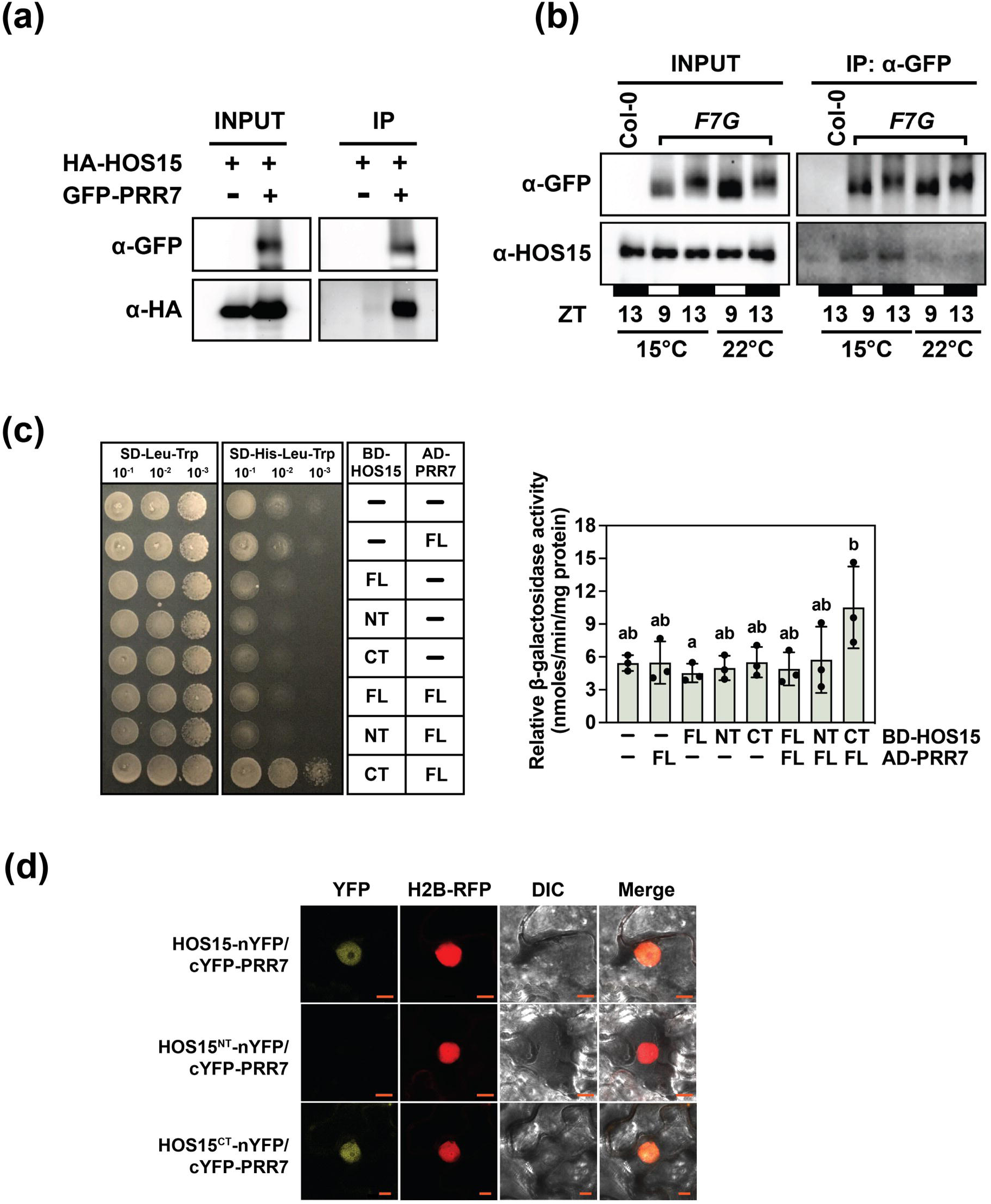
HOS15-PRR7 interaction is enhanced at low temperature. (a) Coimmunoprecipitation of HOS15 and PRR7 in vivo. HA-HOS15 and GFP-PRR7 were transiently expressed in three-week-old N. benthamiana leaves and total proteins were immunoprecipitated with anti- GFP antibody. Eluted immunocomplexes were probed with anti-GFP and anti-HA antibody. Five independent trials were performed. (b) Coimmunoprecipitation of HOS15 and PRR7 in transgenic Arabidopsis plants. F7G was grown under 12L/12D at 22°C for 11 d and transferred to 22°C or 15°C for 3 d. Protein extracts collected at the indicated time points were immunoprecipitated with anti-GFP. Immunodetection performed with with anti-GFP and anti-HOS15 antibody as in Fig 1. Representative of three independent trials is shown. (c) Yeast two-hybrid assay interaction test for HOS15 and PRR7. *HOS15* gene fragments corresponding to the full length (1-613 aa), N-terminal region (1-248 aa; harboring the LiSH domain), and C-terminal region (249-613 aa; harboring the WD40 domain) were fused to the GAL4 binding domain (BD) and co-transformed into AH109 containing the HIS3 genes under the control of GAL4 binding sites, with PRR7 tagged to the GAL4 activation domain (AD). Images were taken after 7 d of incubation of the yeast cells grown on a synthetic SD medium lacking leucine, tryptophan, and histidine at 30°C. β-Galactosidase assay was performed with three independent trials and the mean (nmoles/min/mg protein) ± standard deviation is shown. Letters above bars indicate statistical difference between transformants based on ANOVA and Tukey HSD test. (d) Bimolecular fluorescence complementation assay in N. benthamiana leaves. HOS15 and PRR7 tagged with nEYFP and cEYFP, respectively, were expressed in N. benthamiana leaves with H2B-RFP as a nuclear marker and imaged after 3 d. *HOS15* fragments encoding the full-length gene (1-613 aa), the N-terminal region (1-248 aa), and the C-terminal region (249-613 aa) were separately examined with three independent replicates.

HOS15 contains a LiSH motif and a WD40 domain at the N- and C-terminal regions, respectively. Yeast two-hybrid assays (Y2H) show that the C-terminal region (249-613 aa) of HOS15 is responsible for a direct interaction with PRR7 (Fig. 3c). This was confirmed using BiFC, which also shows that the interaction of HOS15 and PRR7 takes place in the nucleus (Fig. 3d).

A proposed role of HOS15 is to function as a component of the DDB1-cullin4 machinery to recruit substrates to the proteasome-dependent pathway (Park *et al*., 2018). To determine whether HOS15 is involved in PRR7 ubiquitination we tested the level of ubiquitinated PRR7 in the presence or absence of the proteasome inhibitor MG132. Inhibitor-treated immunoprecipitates probed with anti-Ub antibody showed higher levels of ubiquitinated PRR7 in the WT background, relative to the *hos15-2* background, suggesting that PRR7 ubiquitination at low temperatures in the dark requires HOS15 (Fig. 4; Fig S3).

**Figure 4.**
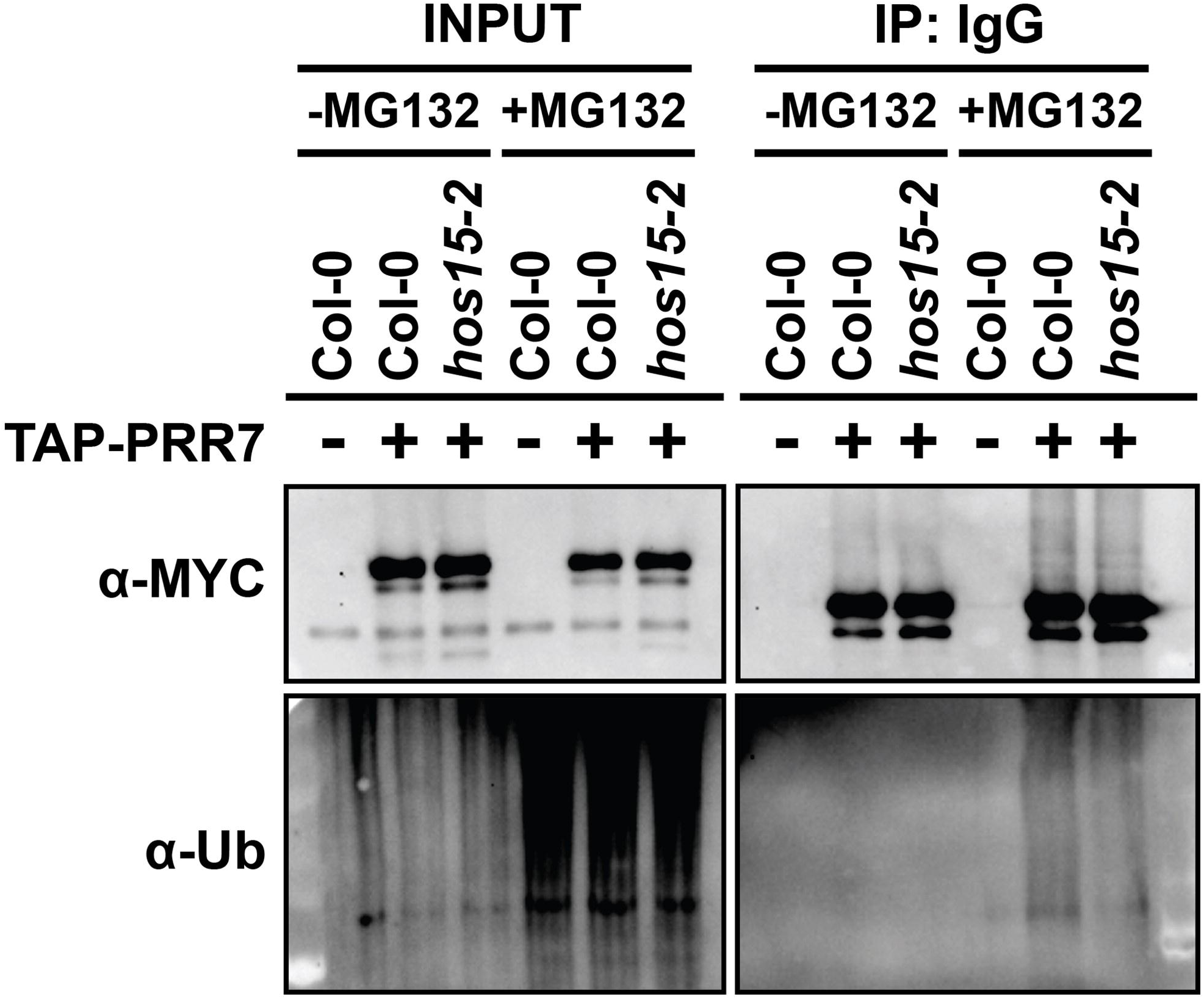
HOS15 mediates PRR7 ubiquitination at low temperature. *In vivo* ubiquitination assay for PRR7. *35S::TAP-PRR7/Col-0* and *35S::TAP-PRR7/hos15-2* plants were grown in 12L/12D at 22°C, followed by 3 d additional 12L/12D at 15°C. The seedlings were collected at ZT13 after incubation with or without MG132 at 15°C. Total proteins were incubated with human IgG-Sepharose and immunocomplexes were probed with anti-myc and anti-Ub antibodies. Representative of two biological replicates.

### *hos15* cold sensitivity requires PRR7

The *hos15-2* mutant is sensitive to freezing stress (Zhu *et al*., 2008; Park *et al*., 2018), whereas *prr7* plants are freezing resistant (Nakamichi *et al*., 2009; Wang *et al*., 2017). To test whether *PRR7* resides within the *HOS15*-mediated cold response pathway, we determined the survival rate of cold-acclimated *hos15-2prr7-3* and WT plants after exposure to a freezing stress protocol, followed by recovery at 15°C (Fig. 5a). The strong freezing sensitivity in the *hos15* mutant is rescued in the *hos15-2prr7-3* mutant to a level comparable to *prr7-3*. Electrolyte leakage assays quantitated these findings (Fig. 5b). Additionally, PRR7 protein abundance in the rosette leaves of *F7G* and *F7Ghos15-2* plants were greatly increased only in the *hos15* mutant background after −4°C and −6°C freezing treatments (Fig. S4), consistent with that observed in seedlings (Fig. 1b). Similarly, plants that were not pre-acclimated at 15°C prior to freezing stress also showed the most severe freezing sensitivity in *hos15-2* (Fig S5). Taken together, these results indicate that PRR7 is necessary for freezing sensitivity and HOS15 is required for reduction of PRR7 levels.

**Figure 5.**
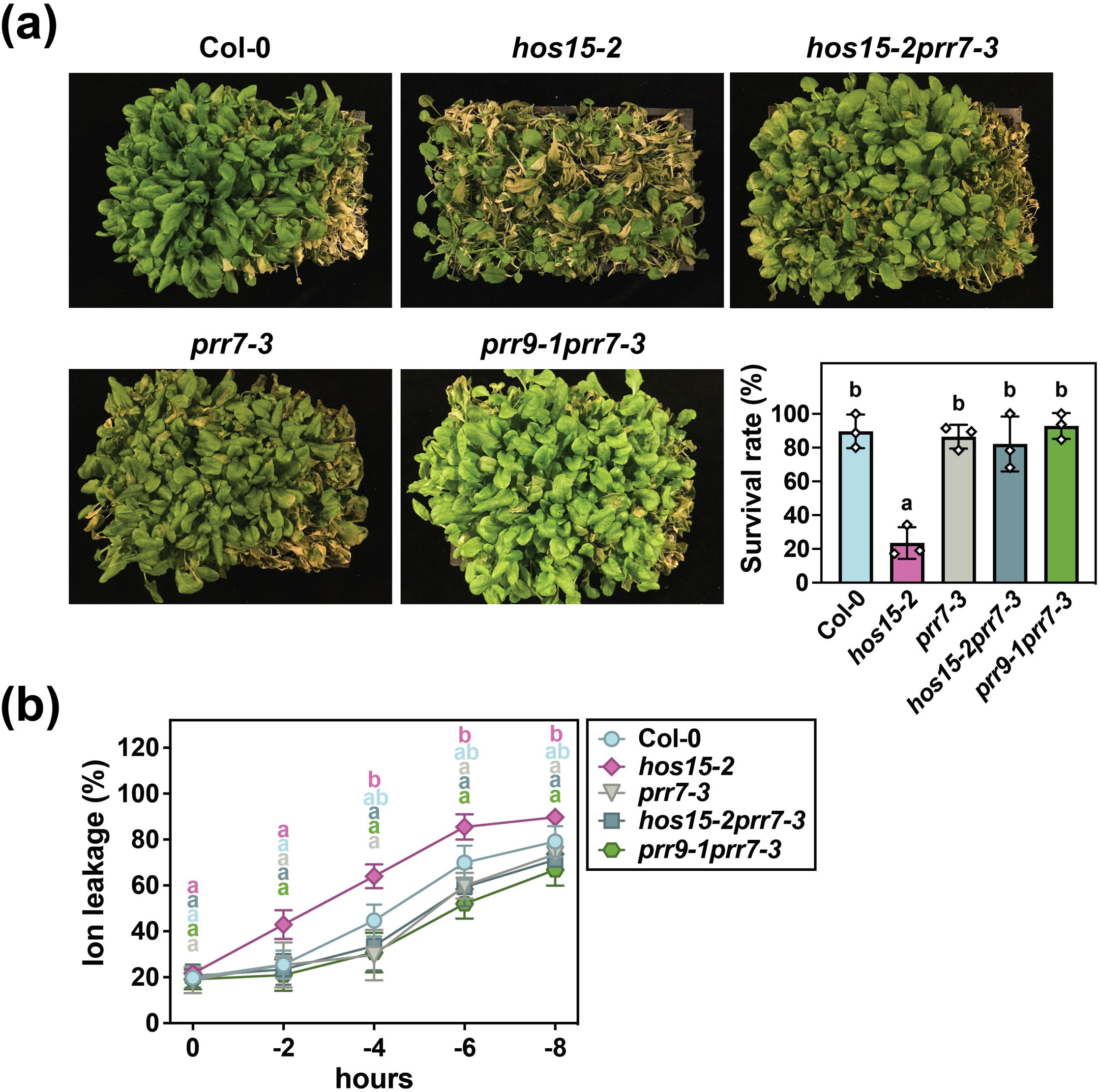
*hos15* cold sensitivity requires PRR7. (a) Effect of *hos15* and *prr7* mutations on plant freezing tolerance. Col-0, *hos15-2*, *prr7-3*, *hos15-2prr7-3*, and *prr9-1prr7-3* were cultivated on soil in 12L/12D at 22°C for 24 d followed by 15°C for one week. A gradual temperature decrease starting at ZT9 h was 4°C for 1 h, 0°C for 1 h, −2°C for 1 h, −4°C for 1 h, −6°C for 1 h, 4°C for 16 h (all above in darkness), 15°C for 7 d (in 12L/12D). Three independent experiments were conducted and the mean of the survival rate (100 – yellow plants/green plants) was plotted with error bars depicting the standard deviation. (b) Effect of *hos15* and *prr7* mutations on freezing-induced electrolyte leakage. Rosette leaves were collected during the freezing treatment at the indicated time points. The mean proportion of electrolyte leakage before and after boiling of the collected leaves is shown, with error bars representing the standard deviation obtained from four independent trials. Letters indicate the significant difference as determined by ANOVA and Tukey HSD test.

### *CBF1* repression by PRR7 is relieved by HOS15

Downstream regulators of the *PRR7*/*HOS15* cold response pathway include the *CBFs*, whose promoters can be occupied by PRR7 (Nakamichi *et al*., 2012; Liu *et al*., 2013; Liu *et al*., 2015) and HOS15 (Park *et al*., 2018). To determine whether *PRR7* and *HOS15* affect the transcriptional abundance of the *CBFs* at 15°C in the dark, we measured the transcript levels of *CBF1*, *CBF2*, and *CBF3* under warm and cold temperatures in four different genetic backgrounds (Fig. 6, Table S1). The expression waveforms of the three genes are strikingly different between the two temperatures. Notably, at 15°C *CBF1* expression remains depressed at ZT13 in *hos15* while levels are more than 3 fold higher in the WT, *prr7-3* and *hos15-2prr7-3* backgrounds. Such a disparity is not observed at 22°C (Fig 6a and b). In contrast. *CBF2* expression over the time course is very similar among all genotypes at 15°C, but the absence of PRR7 at 22°C during the day nearly doubles *CBF2* expression at ZT 5, 9 and 13 (Fig 6a and b). There are no significant differences among the genotypes in *CBF3* levels at either temperature (Fig 6; Table S1). Taken together, these results support a role for PRR7 in the repression of *CBF1* expression at ZT 13 in the dark and cold, which is relieved in the absence of PRR7. Higher levels of PRR7 in the *hos15* background (Fig 1b) are consistent with greater repression of CBF1 at ZT13 (Fig 6b). In contrast, at 22°C, PRR7 levels during the day repress *CBF2* independent of the presence or absence of HOS15. Experiments conducted with much older, soil-grown plants (Fig. 5; Fig. S4), showed very similar reductions in CBF1 and CBF2 levels in the hos15 background (Fig. S6), emphasizing the importance of HOS15-mediated regulation of PRR7 at all stages of development,

**Figure 6.**
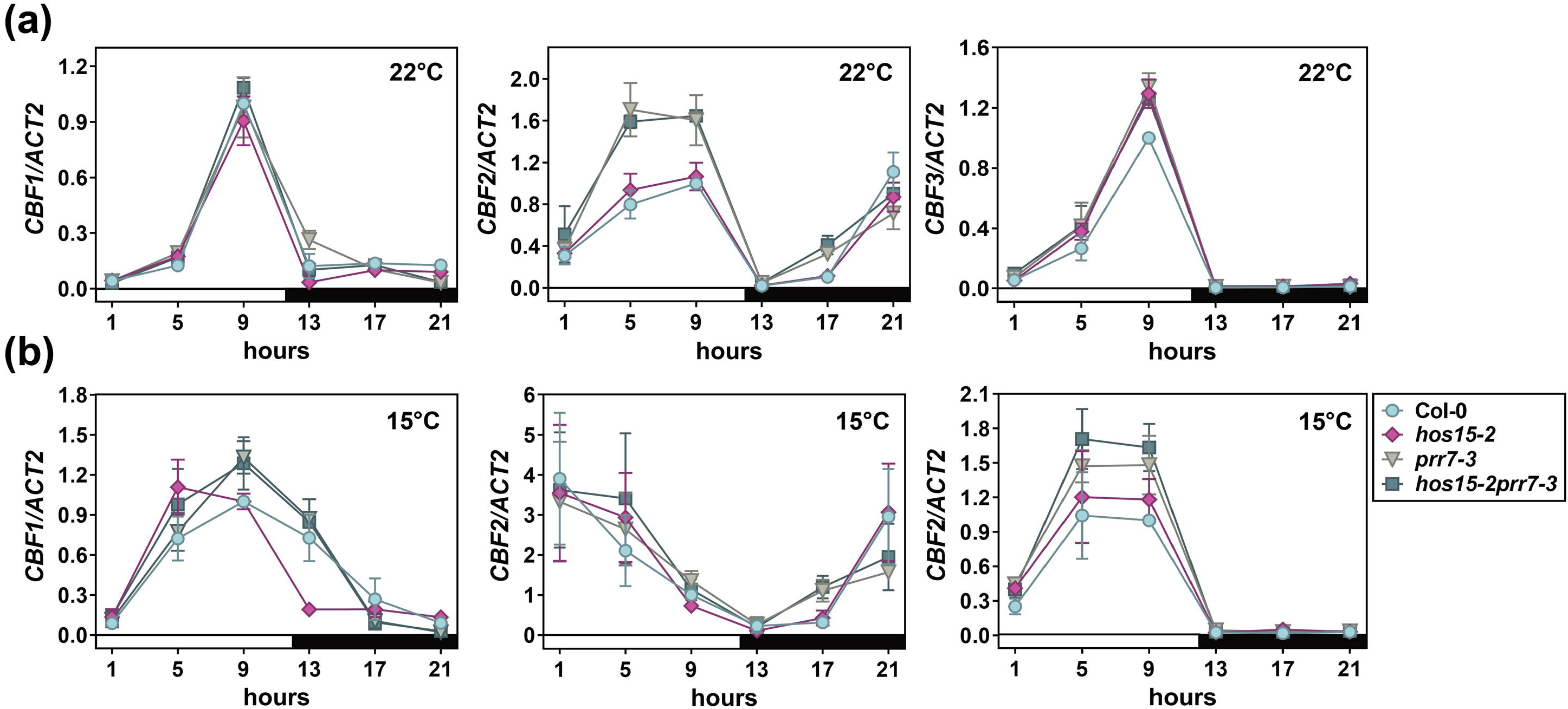
*CBF1* repression by PRR7 is relieved by HOS15 at low night temperatures. Col-0, *hos15-2*, *prr7-3,* and *hos15-2prr7-3* were grown at 22°C for 7 d followed by 22°C (a) or 15°C (b) for 3 d under 12L/12D with tissue collected every 4 h starting 1 h after dawn. Transcript levels of *CBF1*, *CBF2*, and *CBF3* were analyzed by RT-qPCR, and values were normalized to *ACTIN2* (*ACT2*; *AT3G18780*) and scaled to that of Col-0 at ZT9. Mean ± standard deviation (n = 3) is shown. Statistical differences were determined by ANOVA and Tukey HSD test and are shown in Table S1. White and dark bars represent light and dark period. The primers used for the RT-qPCR are listed in Table S2.

### HOS15 decreases levels of PRR7 at *CBF* promoters

We next determined whether HOS15 affects PRR7 residency at the *CBF* promoters, focusing on ZT9 and ZT13 at 15°C. As previous work showed PRR7 enrichment at G-box motifs (Nakamichi *et al*., 2012; Liu *et al*., 2013; Liu *et al*., 2015), we performed ChIP-qPCR, targeting the G-box motifs of the *CBF* promoter regions and the previously identified PRR7 binding regions (Liu *et al*., 2015) (Fig. 7a). PRR7 associates with most regions but is particularly enriched in the *hos15-2* background at ZT13 at the G-box regions at the promoters of *CBF1* and *CBF2* (C and G regions, respectively) (Fig. 7b, d). A significant difference in the enrichment of PRR7 was also observed for other regions, such as the PRR7-binding region B (*CBF1*) (Fig. 7b). Taken together these results indicate that HOS15 functions to decrease the enrichment of PRR7 on the *CBFs* promoters, consistent with higher levels of PRR7 in *hos15* under cold and dark conditions (Fig. 1b).

**Figure 7.**
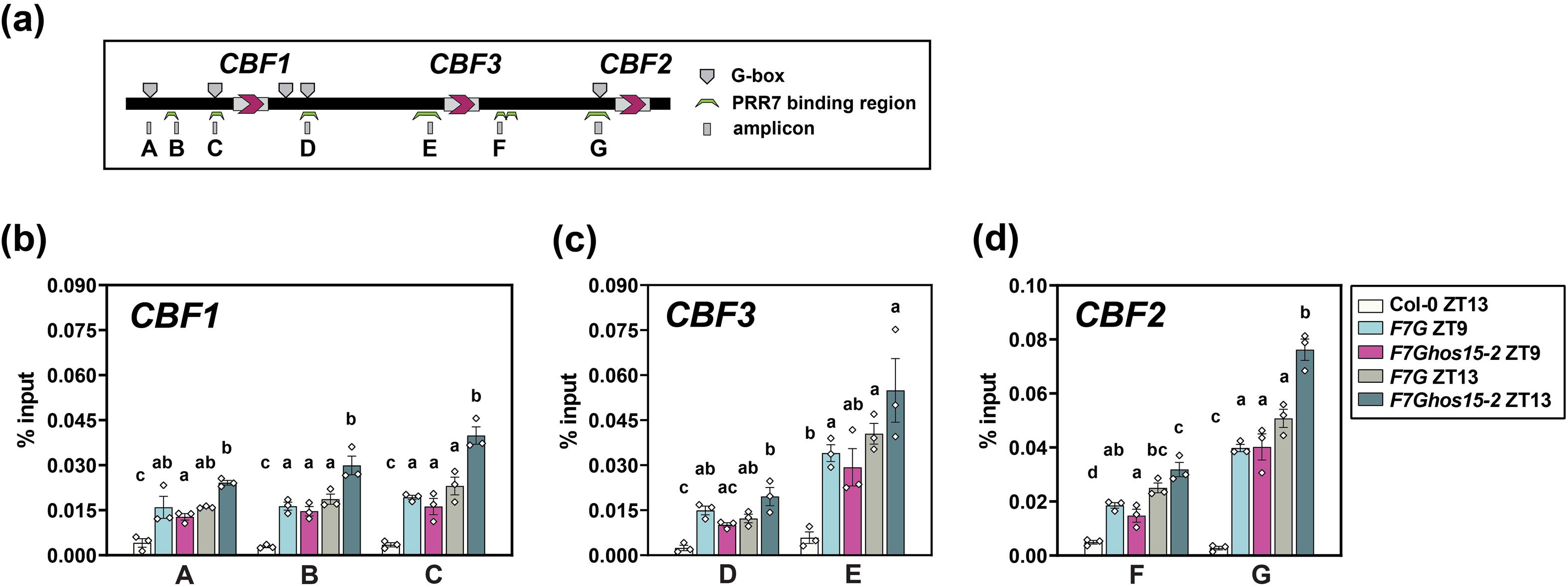
*HOS15* decreases PRR7 enrichment at CBF1/2/3 promoters at low night temperatures. (A) ChIP-qPCR primer binding sites against *CBF* promoter regions. The positions of the G-box motif (CACGTG) and previously identified PRR7 binding sites (Liu *et al*., 2013) and the regions of the amplicons targeted by the ChIP-qPCR binding primers are indicated. (B,C,D) The promoter occupancy of PRR7 on the *CBF1* (A)*, CBF3* (B), and *CBF2* (C) promoter regions. Col-0, *F7G* and *F7Ghos15-2* were grown in 12L/12D at 22°C for 10 d followed by growth at 15°C for 3 d. Seedlings were collected at ZT9 and/or ZT13, and nuclei were isolated. DNA-protein complexes were immunoprecipitated with anti-GFP and the resulting elutes were analyzed by RT-qPCR using the primers targeting the 5’ UTR, 3’ UTR, and gene coding regions of *CBF1, CBF2,* and *CBF3* (Jiang *et al*., 2017). Mean ± standard error (*n* = 3) is shown. The primers used in this assay are in Table S2.

### PRR7 repression of *COR15A* is HOS15-dependent

Directly downstream of the *CBF*s in the regulation of cold tolerance are the cold regulated (*COR*) genes, which are activated in response to CBF binding to *cold and dehydration regulatory elements* (CRT/DRE) in the *COR* promoters (Shi *et al*., 2018; Hwarari *et al*., 2022). Inspection of the *COR15A* promoter revealed three G-box sequences; a known PRR7 binding motif (Liu *et al*., 2013) (Fig 8a). Through ChIP-qPCR we tested PRR7 presence at ZT9 and ZT13 in the presence or absence of HOS15. Only one (amplicon C; Fig 8a) showed enrichment of PRR7 chromatin residence in the *hos15* background at cold temperatures (15 C) and in the dark (ZT13) (Fig 8b). Consistent with this result, *COR15A* expression is repressed under these conditions, most acutely and significantly at ZT13 in the *hos15* background, relative to WT (Fig 8c), but not at 22 C (Fig. 8b). Notably, PRR7 is important in the repression of *COR15A* at ZT9 in the light at both temperatures, and also at ZT13 at 22 C and ZT 5 at 15 C, but all of these effects are independent of the presence of HOS15 (Fig 8b, c; Table S1), suggesting a different mechanism of *COR15A* regulation by PRR7 under these conditions. Taken together, these findings support the notion that under dark and cold conditions PRR7 is also directly repressing *COR15A* gene expression in a HOS15-dependent way, in addition to its role in *CBF1* repression. This additional target in the ICE-CBF-COR cold tolerance pathway enhances the role of PRR7 in the circadian control of cold acclimation (Fig. 9).

**Figure 8.**
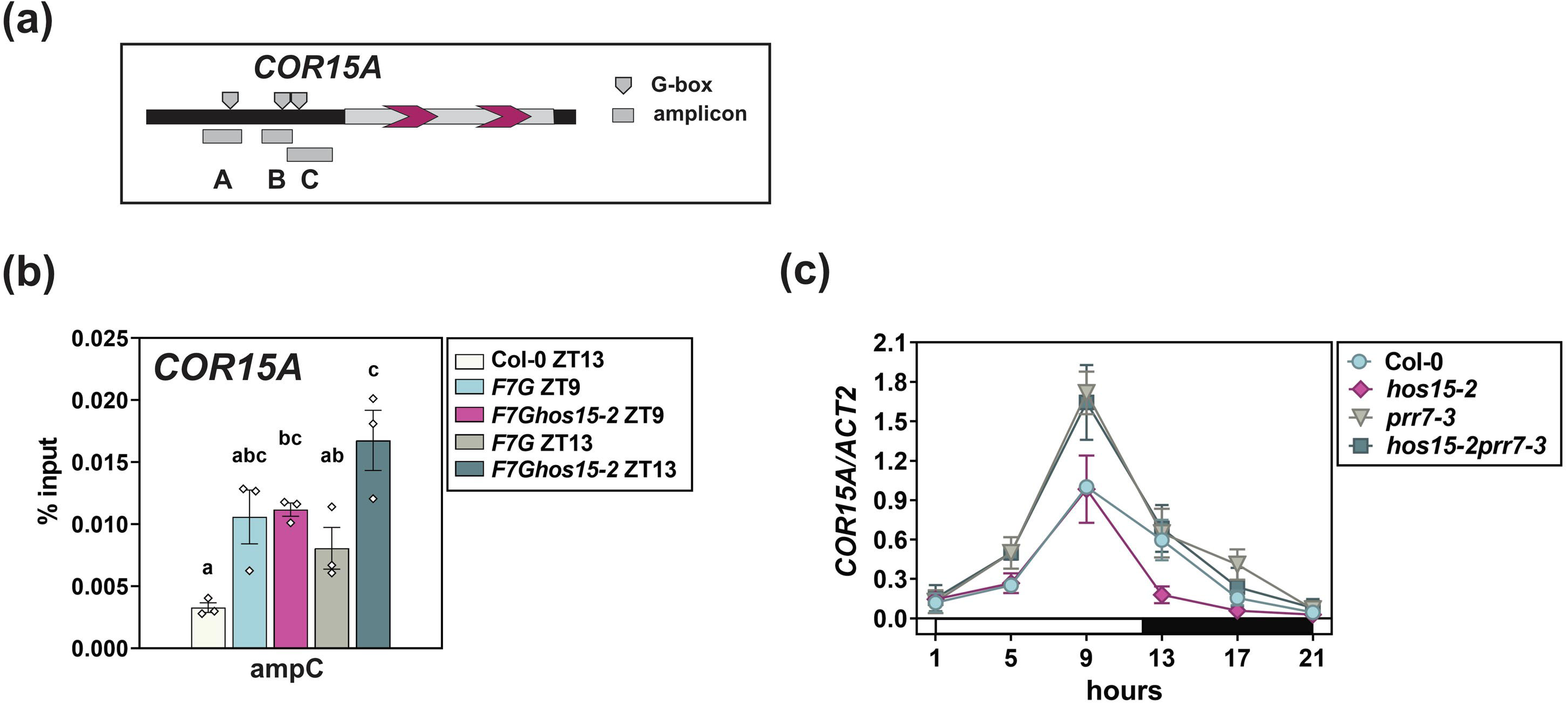
*HOS15* suppression of PRR7 enrichment at the *COR15A* promoter at low night temperatures increases *COR15A* expression. (a) Potential PRR7 binding sites at the *COR15A* promoter. The positions of the G-box motif (CACGTG) and the primers used to test for PRR7 residence via ChIP-qPCR are shown. (b) Chromatin occupancy of PRR7 at amplicon C (C) on the *COR15A* promoter. Col-0, *F7G,* and *F7Ghos15-2* were grown in 12L/12D at 22°C for 10 d followed by growth at 15°C for 3 d. Seedlings were collected at ZT9 and ZT13, and DNA-protein complexes were immunoprecipitated with anti-GFP then analyzed by RT-qPCR. Mean ± standard error (*n* = 3) is shown. Primers listed in Table S2. (c, d) *COR15A* expression in seedlings grown in 12L/12D at 22°C for 10 d followed by growth at 22°C (c) or 15°C (d) for 3 d. Statistical differences determined by ANOVA and the Tukey HSD test are shown in Table S1. Mean ± standard deviation (n=3) is shown.

**Figure 9.**
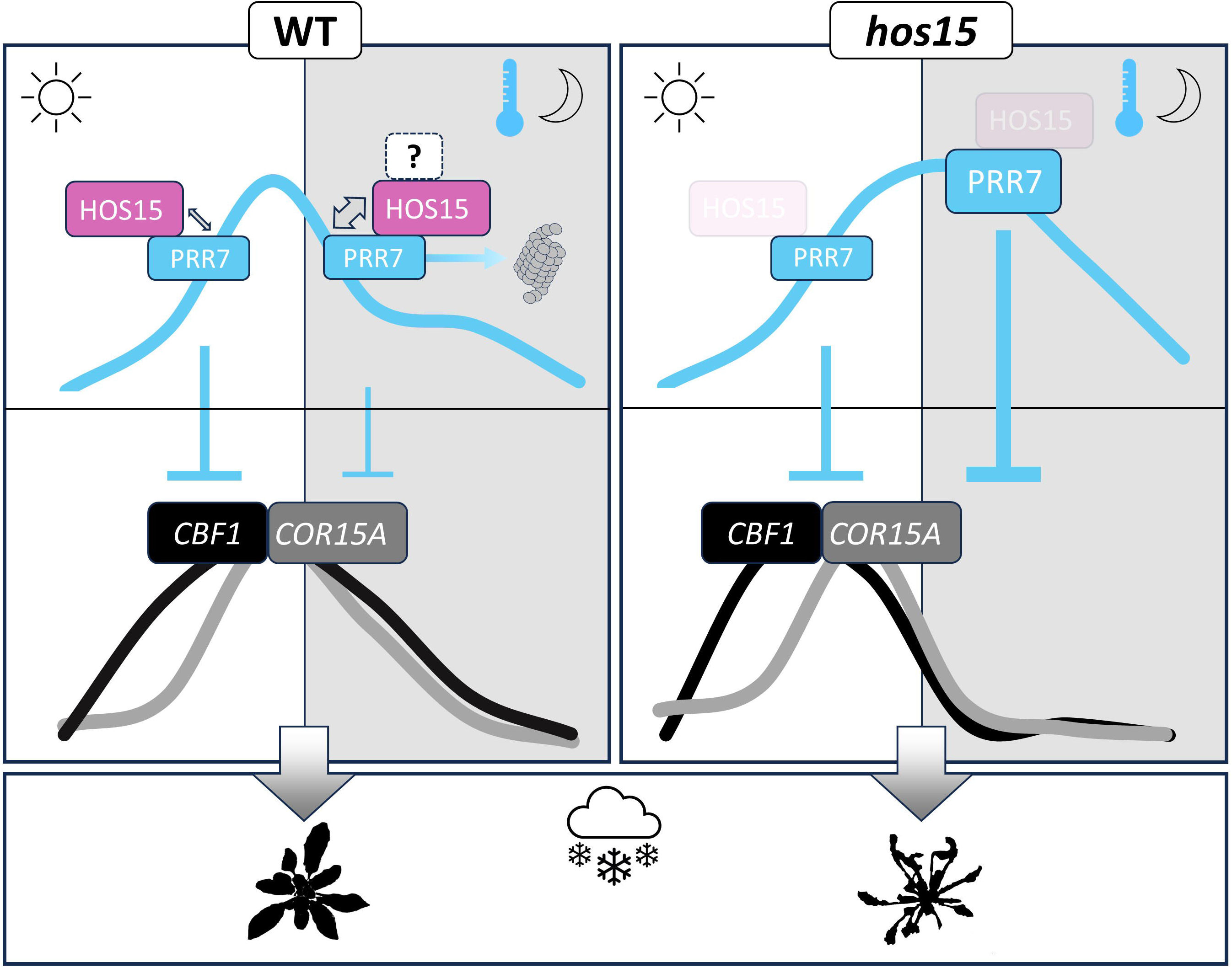
A working model of the post-translational regulation of PRR7 abundance and enhanced cold tolerance by HOS15. HOS15 promotes the degradation of the PRR7 protein through a CULLIN4-based E3 ubiquitin ligase complex (proteasome icon) during cold nights, resulting in the transcriptional de-repression of the cold stress-responsive genes *CBF1* and *COR15A*. In the *hos15* mutant PRR7 levels remain high at night, *CBF1* and *COR15A* levels are repressed, resulting in increased freezing sensitivity (shriveled silhouette in lower panel). **?**: indicates unknown factors that facilitate the stronger HOS15-PRR7 interaction (indicated by a gray double-head arrow) in the dark and cold. Thickness of double headed arrows indicates relative strength of the HOS15-PRR7 interaction.

## Discussion

The PRR family of transcriptional repressors play essential roles both in the maintenance of the circadian oscillator, and in providing phase-specificity to a wide range of molecular outputs. Each of the family members has distinct but overlapping expression profiles, the discreteness made possible by relatively short protein half-lives (Farré & Kay, 2007; Kiba *et al*., 2007; Fujiwara *et al*., 2008; Nakamichi, N. *et al*., 2010; Wang *et al*., 2010; Yan *et al*., 2021). The factors responsible for the turnover of TOC1 and PRR5 have been identified (Más *et al*., 2003; Kiba *et al*., 2007) but for most other family members it remains unclear.

Our results now establish that HOS15 promotes the PRR7 protein turnover under very specific conditions, namely, at moderately low temperatures in darkness, likely by recruiting the PRR7 to an ubiquitin E3 ligase complex comprised in part by HOS15. This finding significantly expands the role of HOS15 in the control of plant responses to low temperature (Fig 9).

Previously, HOS15 has been described as a component of a DDB1-Cullin4 complex to target substrates for proteasome-dependent degradation (Park *et al*., 2018). These targets have included HD2C and GI, with the effect most pronounced at low temperatures (Park *et al*., 2018; Ahn *et al*., 2023). In our stability assays, the rate of PRR7 decrease at 22°C is very similar in the WT and *hos15* backgrounds regardless of light conditions (Fig. 2a,b). In contrast, under darkness at 15°C, a decrease in the PRR7 turnover rate is evident in the *hos15* mutant, relative to WT (Fig. 2d). This result aligns with previous work, and now demonstrates a broadened role for HOS15 in low temperature proteolysis.

A current model for freezing tolerance puts HOS15 in association with a protein complex consisting of POWERDRESS (PWR) and HD2C to modulate chromatin modification of cold-regulated (*CORs*) genes (Park *et al*., 2018; Lim *et al*., 2020). Degradation of HD2C by HOS15 at low temperatures induces an increase in H3 acetylation at *COR* promoters resulting in increased CBF occupancy, an up-regulation of *COR* genes and enhanced cold tolerance (Park *et al*., 2018; Lim *et al*., 2020). Our results now indicate that HOS15 can also promote cold tolerance through the alleviation of *CBF1* and *COR15A* repression by facilitating PRR7 turnover (Fig. 2d, 4,6,7,8), supporting previous studies implicating the *CBF*s as possible targets of PRR7 (Nakamichi *et al*., 2009; Liu *et al*., 2013; Liu *et al*., 2015). This additional proteolytic target of HOS15 multiplies its effect by removing an additional repressor of cold acclimation, namely PRR7, and directly connects its function to the circadian control of freezing tolerance.

Interestingly, expression of both *PRR7* and the *CBF*s are clock controlled (Dong *et al*., 2011) and PRR7 would normally represses *CBF1* at dusk to alter its waveform (Fig 6b; *hos15* background), but for the action of the constitutively expressed HOS15 (Ahn *et al*., 2023) altering the PRR7 waveform under cold temperatures (Fig. 1). The same is true for the *COR15A* waveform (Fig 8c) which is similarly dependent on the effect of HOS15 on PRR7 levels.

Notably, only *CBF1* showed a strong *hos15* -dependent drop in expression at low temperatures under our conditions (Fig 6b). Earlier work has shown that *CBF1* loss alone is sufficient to compromise cold tolerance (Novillo *et al*., 2007), suggesting that the phasing and expression level of PRR7 accumulation and its effect on *CBF1* expression is central to the process of cold acclimation. Additionally, CBF1 directly controls *COR15A* (Novillo *et al*., 2007; Wang & Hua, 2009), a second target of PRR7 (Fig 8, 9)(Hwarari *et al*., 2022). COR15A is an intrinsically disordered cold regulated chloroplast protein that contributes to freezing tolerance in Arabidopsis. Overexpression of *COR15A* increases cold tolerance in non-acclimated lines, while knockdown of *COR15A*, and close relative *COR15b*, significantly impairs freezing tolerance of cold-acclimated leaves (Artus *et al*., 1996; Thalhammer *et al*., 2014). Hence, PRR7 has a role at two distinct, but sequential steps in the complex ICE-CBF-COR cold tolerance pathway in Arabidopsis (Fig. 9)(Knight & Knight, 2012; Hwarari *et al*., 2022).

The low-temperature specificity of HOS15 in regulating PRR7 stability can be explained largely by the increased protein-protein interaction at 15°C (Fig. 3b) (Ahn *et al*., 2023). While it remains unknown how low temperature and darkness lead to this increased interaction, a possible mechanism may involve COP1, whose complex formation and the subcellular condensation are regulated in a light- and temperature-dependent manner (Jang *et al*., 2015; Ahn *et al*., 2023). HOS15 interacts with COP1 through its N-terminal region containing a LisH domain and an F-box-like motif (Ahn *et al*., 2023), whereas HOS15 binding to PRR7 occurs through the WD40 domain (Fig. 3c). In transient expression assays in *N. benthamiana*, COP1 interacts with PRR7 (Fig. S7). Previous studies showed that total COP1 protein abundance increases in response to 16°C (Jang *et al*., 2015), and the nuclear and cytoplasmic COP1 appear to be more soluble at 16°C, as manifested by less accumulation in speckles at 16°C compared to 23°C (Ahn *et al*., 2023). In addition to temperature regulation, darkness induces an activation of the COP1/SPA activity (Hoecker, 2017). These results suggest COP1 as a possible candidate that conditionally leads to HOS15 regulation of PRR7 and would be a fruitful direction for further testing.

Previous work has shown that PRR7 and PRR9 work together in sustaining circadian rhythms and contribute together to affect freezing tolerance (Salome & McClung, 2005; Wang *et al*., 2017). Our freezing test assays showed a similar degree of cold tolerance in the *prr7* and *prr7prr9* double mutants (Fig. 5, S5). PRR7 and PRR9 may heterodimerize which could explain why the two mutant phenotypes are not additive (Fig 5). However, at the most severe freezing conditions (−8C; Fig S5) the *prr7prr9* mutant was slightly more resistant than the *hos15prr7* plants. This type of result would be expected if HOS15 also controls PRR9 levels. Future tests the degree of overlap and independence of both closely related genes in the context of cold acclimation would likely be illuminating to the field.

The factors and signaling pathways involved in cold acclimation and tolerance are very complex and multi-factorial (Shi et al., 2018; Ding & Yang, 2022). For example, all mutant backgrounds, including *hos15,* show very high levels of cold temperature induction of *CBF* expression after only 4 hours of freezing stress (Fig S6), demonstrating a significant degree of HOS15-independent control of this key family of cold tolerance genes. Our work here provides new insight into one aspect of the complexity of cold acclimation: how the circadian system intersects with temperature signals to post-translationally alter protein profiles that lead to increased cold tolerance.

## Acknowledgements

This work was supported by the National Institutes of Health (R01GM093285 and R35GM136400 to D.E.S.), the Next-Generation BioGreen21 Program (PJ01327305), the Rural Development Administration, Republic of Korea (to D.E.S.)

## Author contributions

YJK and DS designed the research, analyzed the data, and wrote the manuscript. YJK performed the research. WYK contributed reagents and discussed the manuscript.

## Data availability

The data that support the findings of this study are available in the article and in the Supporting Information of this article.

