## Supplemental Figures for "HOS15-mediated turnover of PRR7 enhances freezing tolerance"

### Slide 1
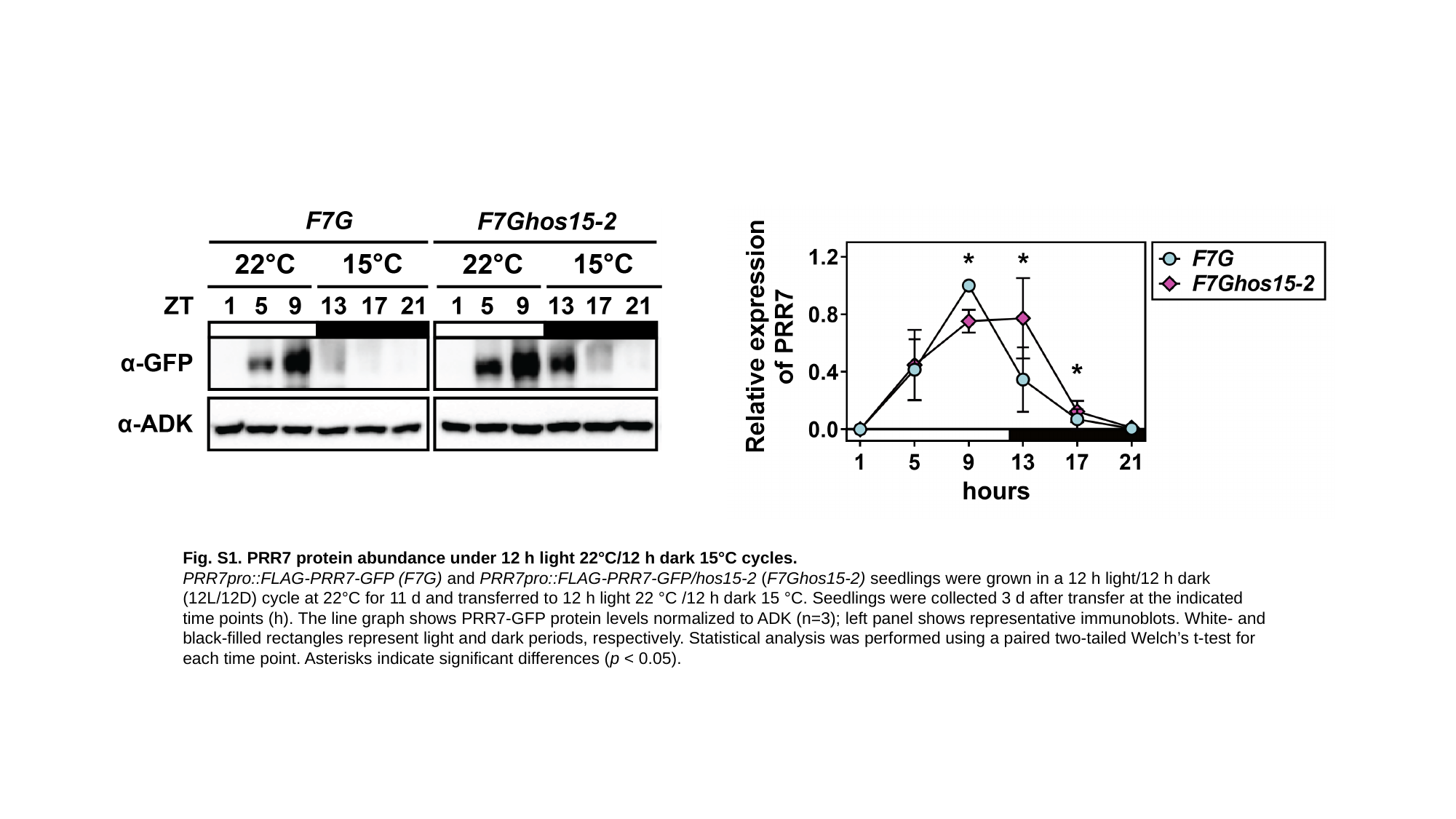

Fig. S1. PRR7 protein abundance under 12 h light 22°C/12 h dark 15°C cycles.
PRR7pro::FLAG-PRR7-GFP (F7G) and PRR7pro::FLAG-PRR7-GFP/hos15-2 (F7Ghos15-2) seedlings were grown in a 12 h light/12 h dark (12L/12D) cycle at 22°C for 11 d and transferred to 12 h light 22 °C /12 h dark 15 °C. Seedlings were collected 3 d after transfer at the indicated time points (h). The line graph shows PRR7-GFP protein levels normalized to ADK (n=3); left panel shows representative immunoblots. White- and black-filled rectangles represent light and dark periods, respectively. Statistical analysis was performed using a paired two-tailed Welch’s t-test for each time point. Asterisks indicate significant differences (p < 0.05).

### Slide 2
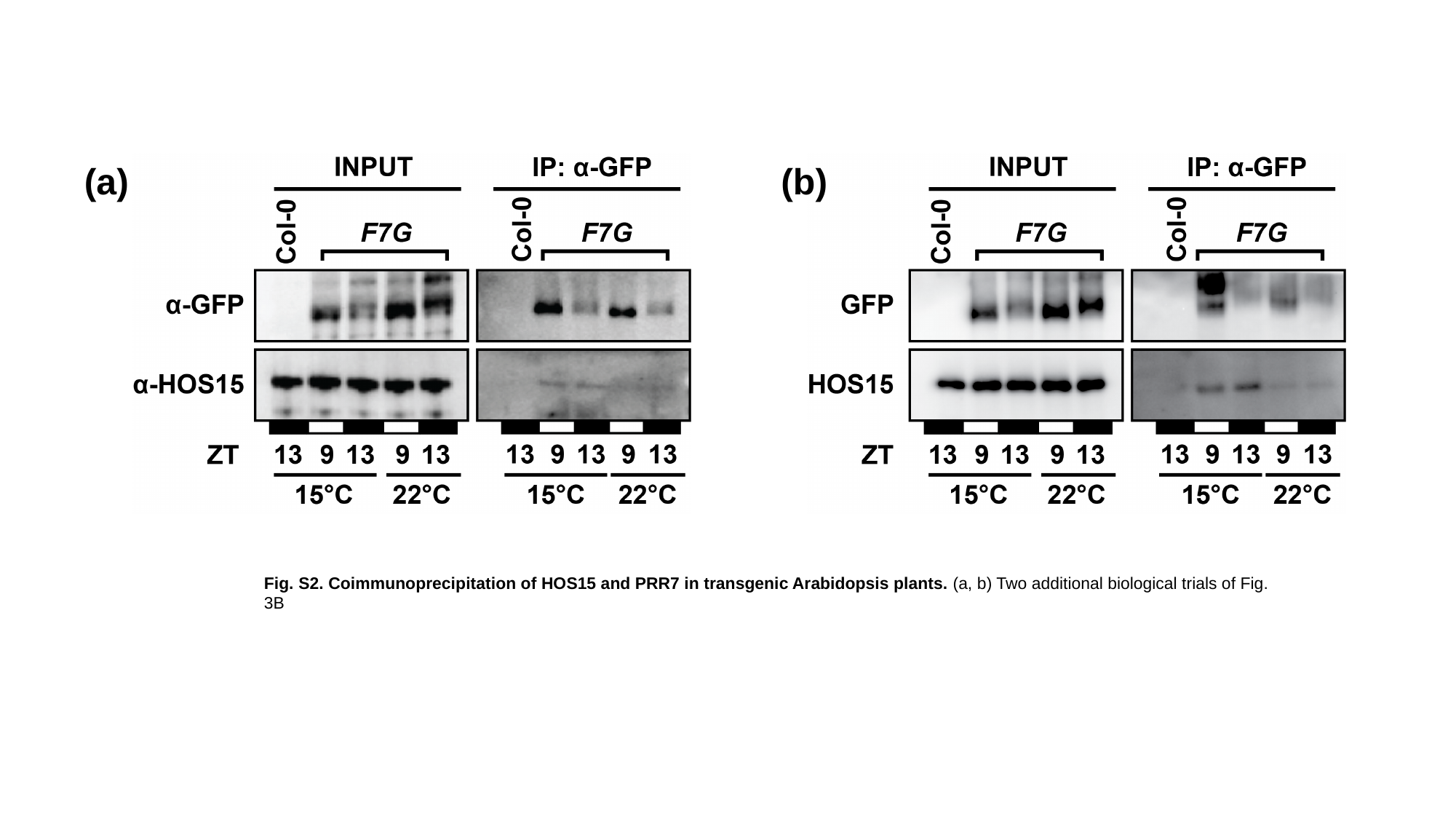

(a)
(b)
Fig. S2. Coimmunoprecipitation of HOS15 and PRR7 in transgenic Arabidopsis plants. (a, b) Two additional biological trials of Fig. 3B

### Slide 3
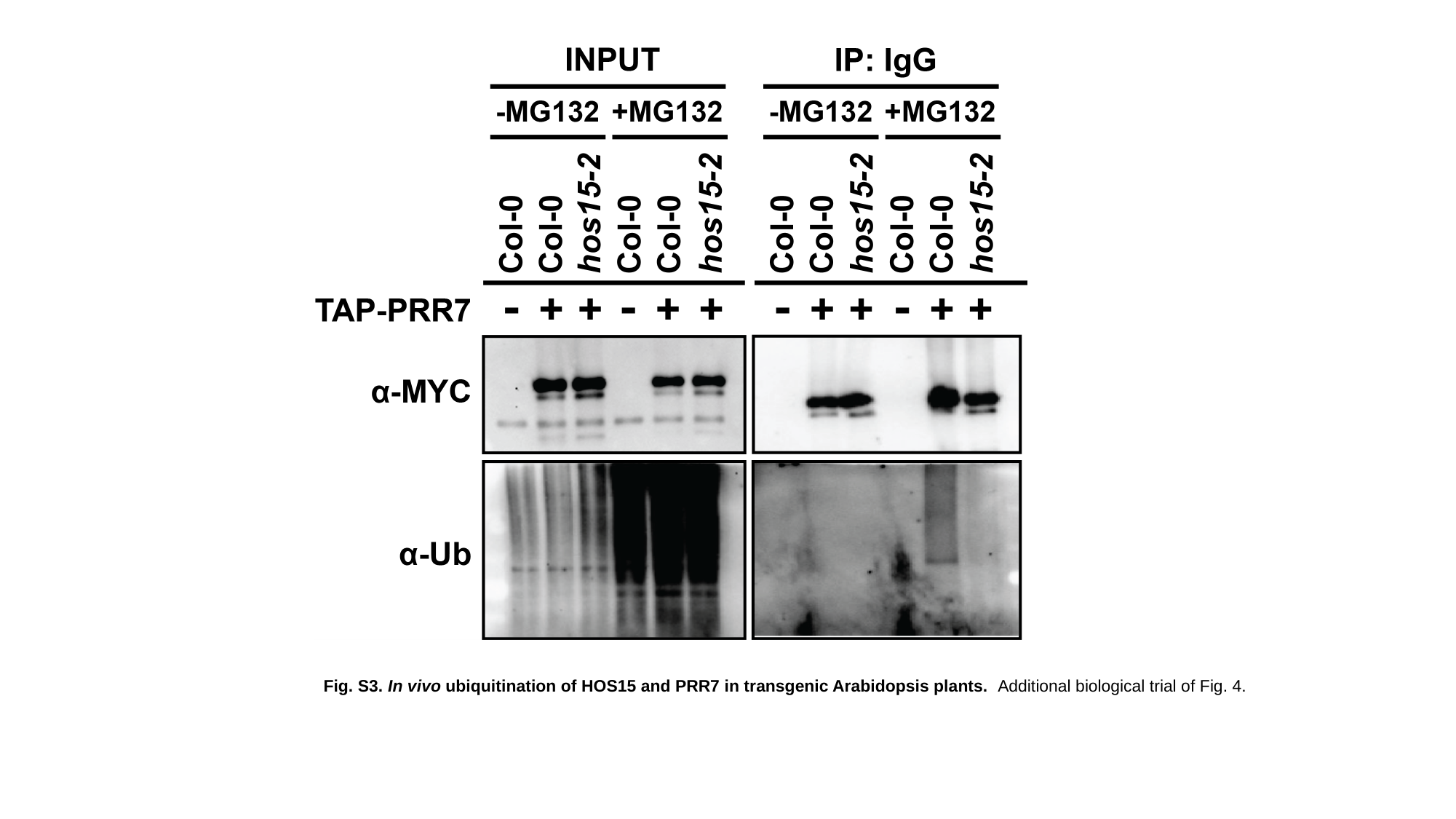

Fig. S3. In vivo ubiquitination of HOS15 and PRR7 in transgenic Arabidopsis plants. Additional biological trial of Fig. 4.

### Slide 4
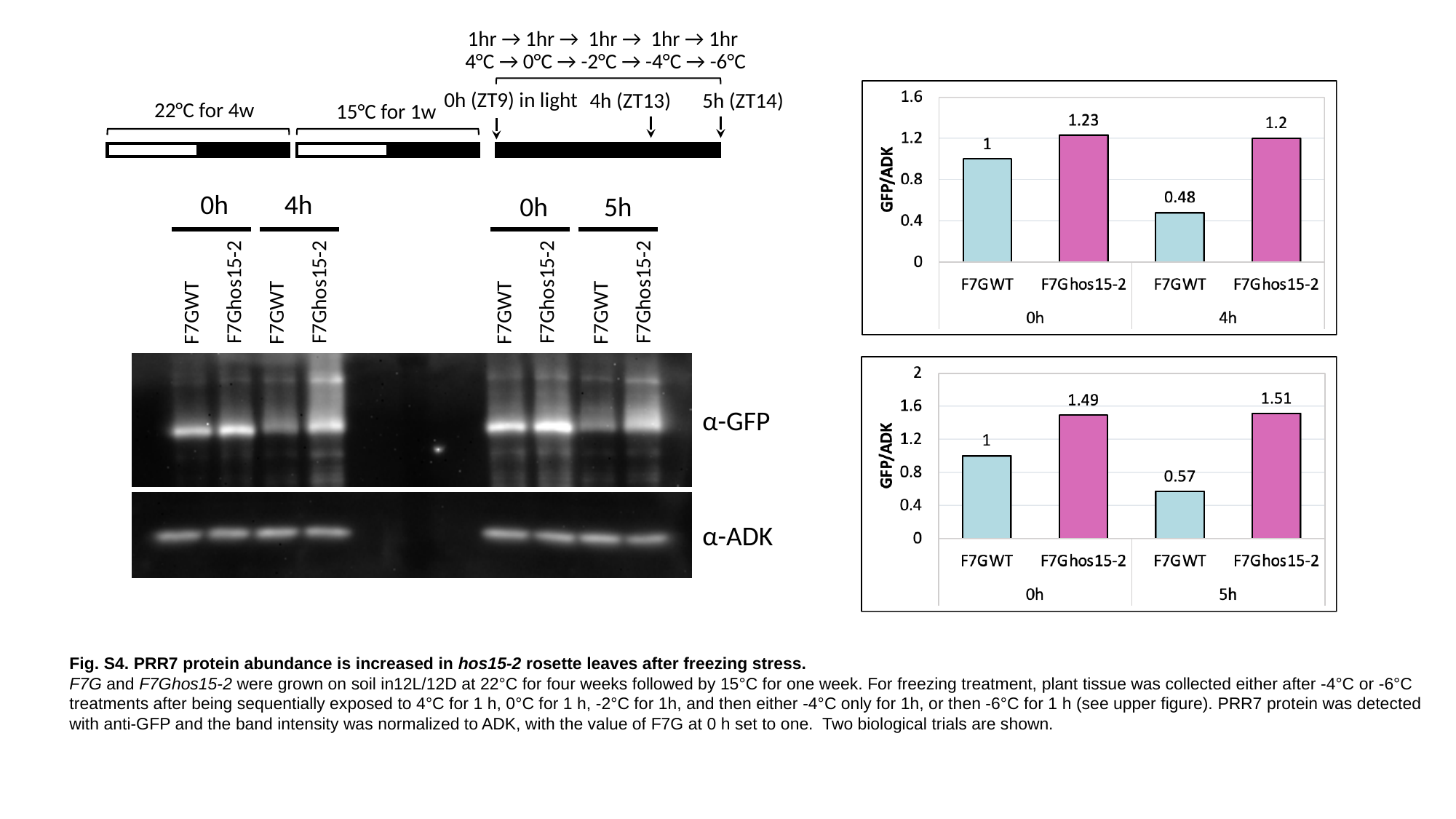

1hr → 1hr → 1hr → 1hr → 1hr
4°C → 0°C → -2°C → -4°C → -6°C
0h (ZT9) in light
4h (ZT13)
5h (ZT14)
22°C for 4w
15°C for 1w
0h
4h
0h
5h
F7Ghos15-2
F7Ghos15-2
F7Ghos15-2
F7Ghos15-2
F7GWT
F7GWT
F7GWT
F7GWT
α-GFP
α-ADK
Fig. S4. PRR7 protein abundance is increased in hos15-2 rosette leaves after freezing stress.
F7G and F7Ghos15-2 were grown on soil in12L/12D at 22°C for four weeks followed by 15°C for one week. For freezing treatment, plant tissue was collected either after -4°C or -6°C treatments after being sequentially exposed to 4°C for 1 h, 0°C for 1 h, -2°C for 1h, and then either -4°C only for 1h, or then -6°C for 1 h (see upper figure). PRR7 protein was detected with anti-GFP and the band intensity was normalized to ADK, with the value of F7G at 0 h set to one. Two biological trials are shown.

### Slide 5
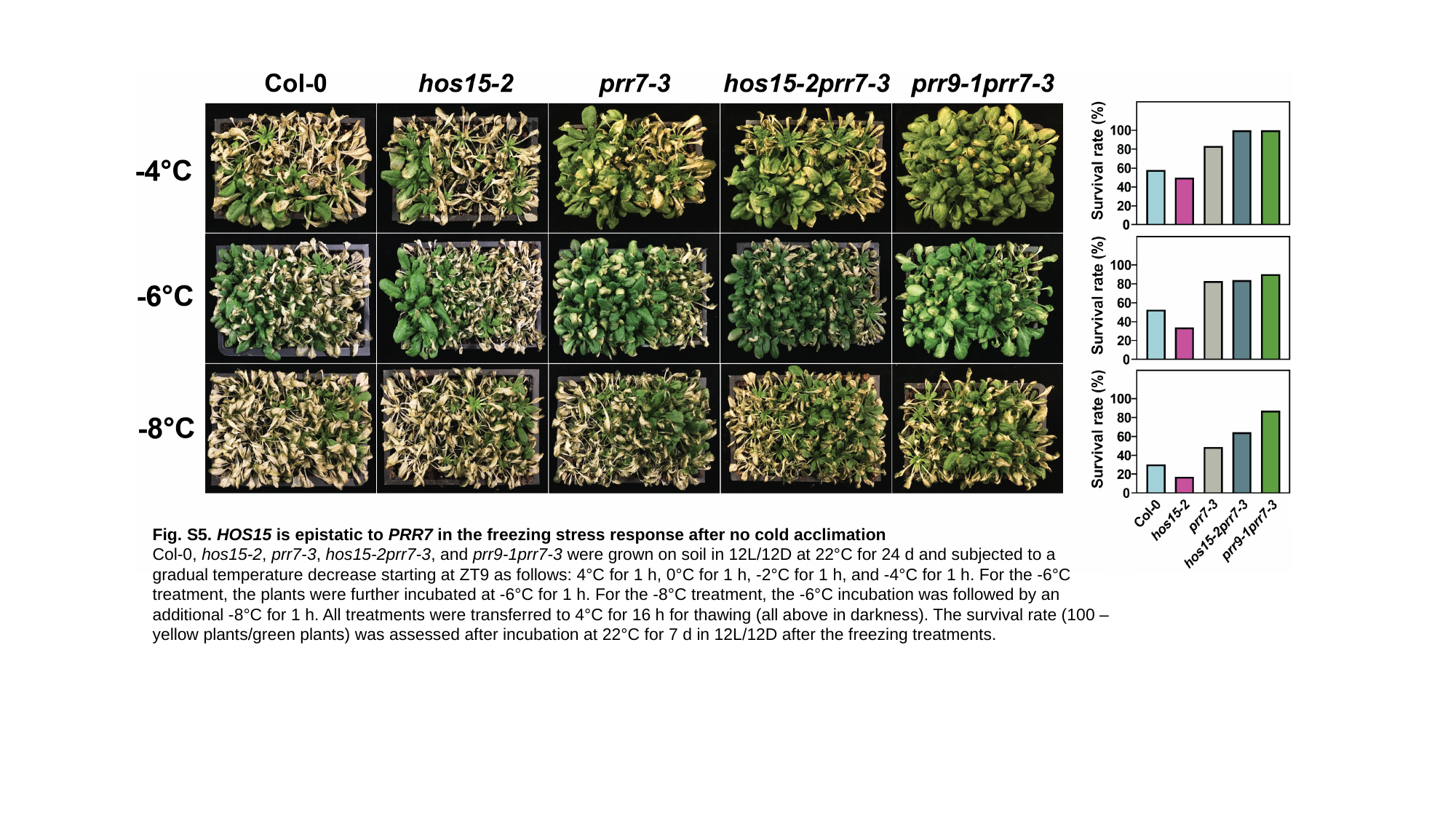

Fig. S5. HOS15 is epistatic to PRR7 in the freezing stress response after no cold acclimation
Col-0, hos15-2, prr7-3, hos15-2prr7-3, and prr9-1prr7-3 were grown on soil in 12L/12D at 22°C for 24 d and subjected to a gradual temperature decrease starting at ZT9 as follows: 4°C for 1 h, 0°C for 1 h, -2°C for 1 h, and -4°C for 1 h. For the -6°C treatment, the plants were further incubated at -6°C for 1 h. For the -8°C treatment, the -6°C incubation was followed by an additional -8°C for 1 h. All treatments were transferred to 4°C for 16 h for thawing (all above in darkness). The survival rate (100 – yellow plants/green plants) was assessed after incubation at 22°C for 7 d in 12L/12D after the freezing treatments.

### Slide 6
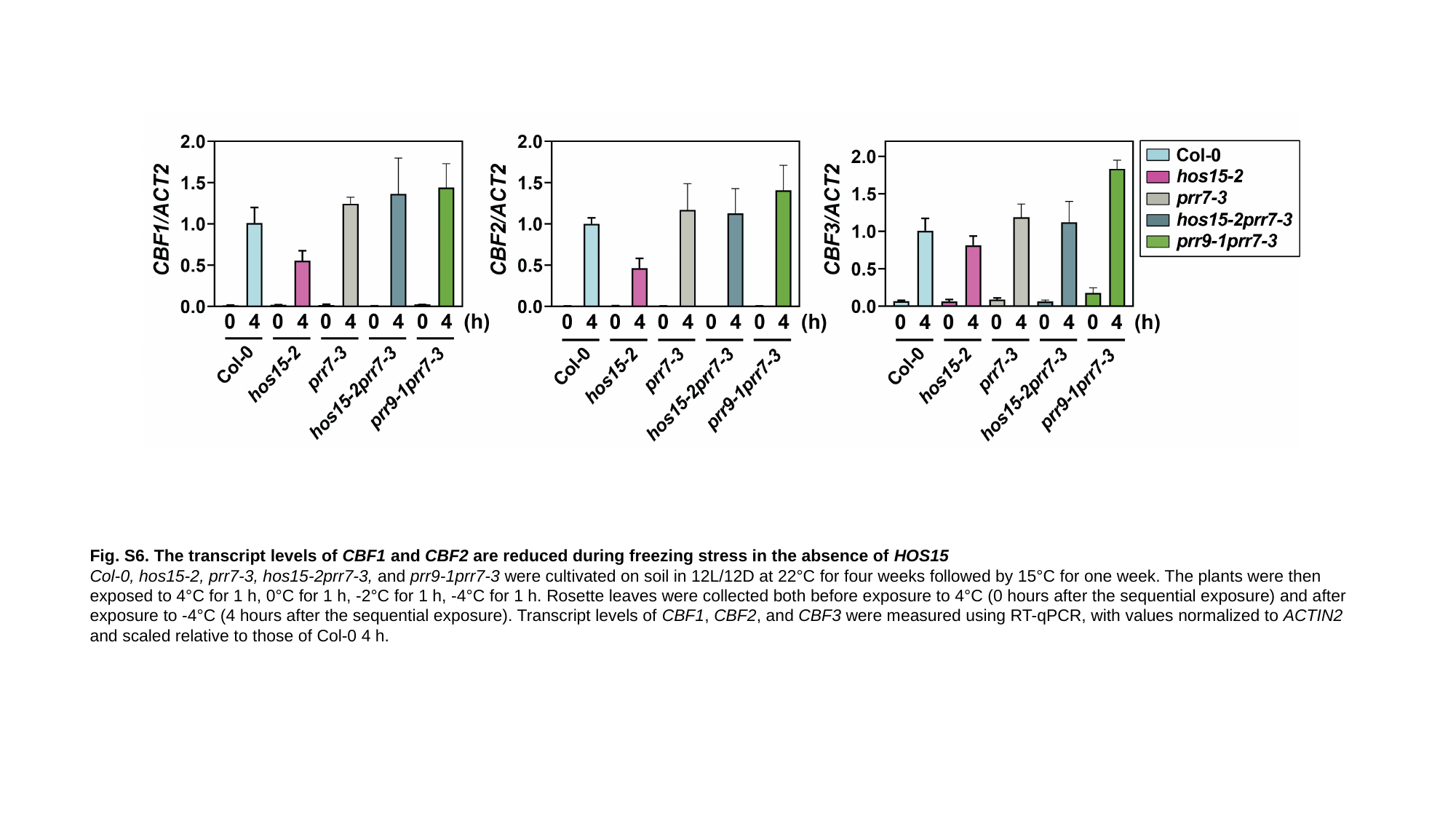

Fig. S6. The transcript levels of CBF1 and CBF2 are reduced during freezing stress in the absence of HOS15
Col-0, hos15-2, prr7-3, hos15-2prr7-3, and prr9-1prr7-3 were cultivated on soil in 12L/12D at 22°C for four weeks followed by 15°C for one week. The plants were then exposed to 4°C for 1 h, 0°C for 1 h, -2°C for 1 h, -4°C for 1 h. Rosette leaves were collected both before exposure to 4°C (0 hours after the sequential exposure) and after exposure to -4°C (4 hours after the sequential exposure). Transcript levels of CBF1, CBF2, and CBF3 were measured using RT-qPCR, with values normalized to ACTIN2 and scaled relative to those of Col-0 4 h.

### Slide 7
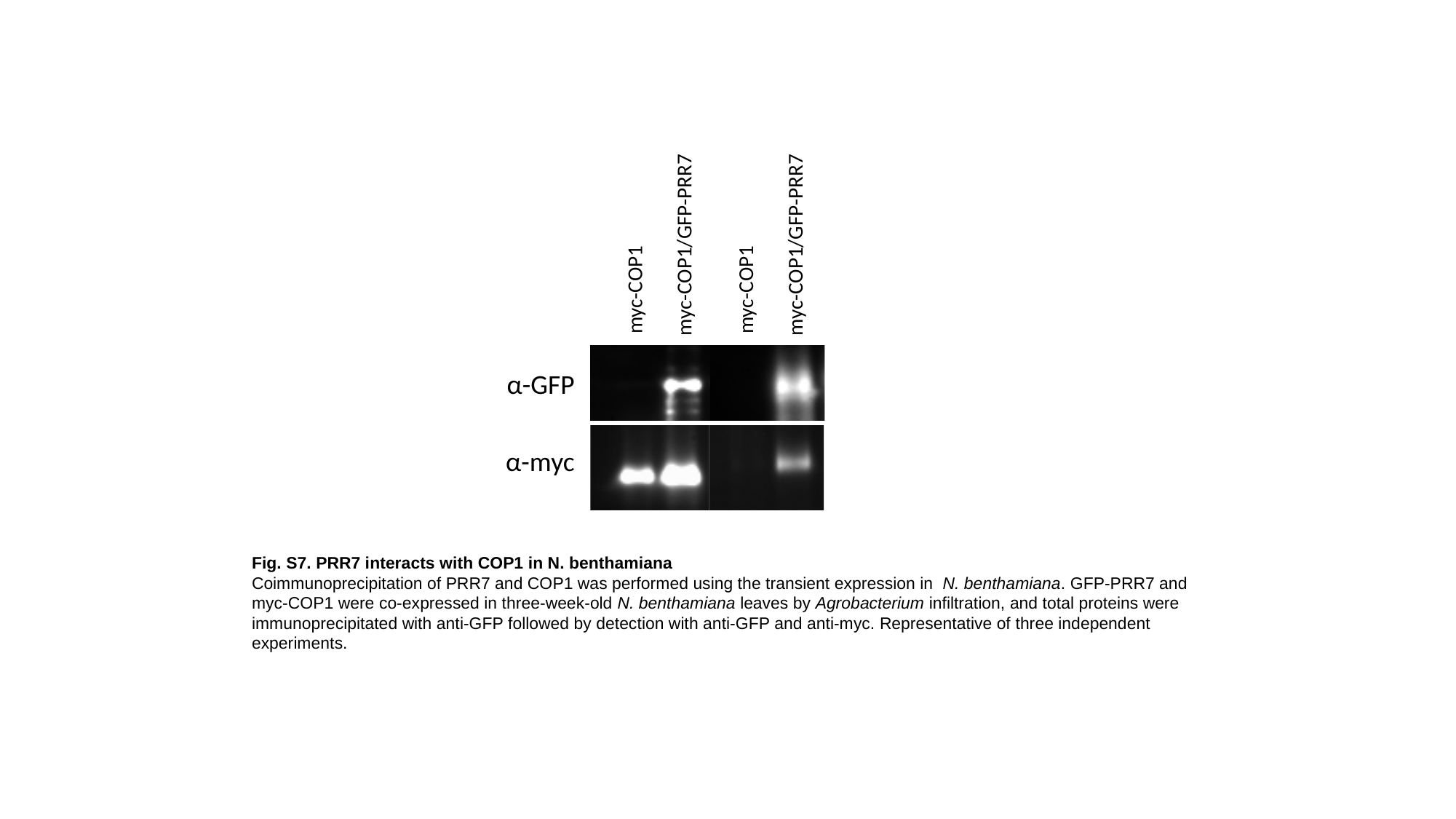

myc-COP1/GFP-PRR7
myc-COP1/GFP-PRR7
myc-COP1
myc-COP1
α-GFP
α-myc
Fig. S7. PRR7 interacts with COP1 in N. benthamiana
Coimmunoprecipitation of PRR7 and COP1 was performed using the transient expression in N. benthamiana. GFP-PRR7 and myc-COP1 were co-expressed in three-week-old N. benthamiana leaves by Agrobacterium infiltration, and total proteins were immunoprecipitated with anti-GFP followed by detection with anti-GFP and anti-myc. Representative of three independent experiments.
