## Supplemental Tables for "HOS15-mediated turnover of PRR7 enhances freezing tolerance"

**Table S1.** **Statistical analysis for RT-qPCR in Figures 6 and 8.**

|  |  | ZT1 | ZT5 | ZT9 | ZT13 | ZT17 | ZT21 |
| --- | --- | --- | --- | --- | --- | --- | --- |
| CBF1 22°C | Col-0 | a | a | a | ab | a | b |
|  | *hos15-2* | a | a | a | a | a | ab |
|  | *prr7-3* | a | a | a | b | a | a |
|  | *hos15-2prr7-3* | a | a | a | ab | a | a |
| CBF2 22°C | Col-0 | a | a | a | a | a | a |
|  | *hos15-2* | a | a | a | a | a | a |
|  | *prr7-3* | a | b | b | a | b | a |
|  | *hos15-2prr7-3* | a | b | b | a | b | a |
| CBF3 22°C | Col-0 | a | a | b | a | b | a |
|  | *hos15-2* | a | a | a | a | ab | a |
|  | *prr7-3* | a | a | a | a | ab | a |
|  | *hos15-2prr7-3* | a | a | a | a | a | a |
| COR15A 22°C | Col-0 | a | a | a | ab | a | ab |
|  | *hos15-2* | a | a | a | a | a | a |
|  | *prr7-3* | b | a | b | b | a | b |
|  | *hos15-2prr7-3* | ab | a | b | b | a | b |
| CBF1 15°C | Col-0 | a | a | a | b | a | ab |
|  | *hos15-2* | a | a | a | a | a | a |
|  | *prr7-3* | a | a | a | b | a | b |
|  | *hos15-2prr7-3* | a | a | a | b | a | b |
| CBF2 15°C | Col-0 | a | a | a | a | b | a |
|  | *hos15-2* | a | a | a | a | ab | a |
|  | *prr7-3* | a | a | a | a | ab | a |
|  | *hos15-2prr7-3* | a | a | a | a | a | a |
| CBF3 15°C | Col-0 | a | a | a | a | a | a |
|  | *hos15-2* | a | a | a | a | a | a |
|  | *prr7-3* | a | a | a | a | a | a |
|  | *hos15-2prr7-3* | a | a | a | a | a | a |
| COR15A 15°C | Col-0 | a | a | a | a | ab | a |
|  | *hos15-2* | a | a | a | b | b | a |
|  | *prr7-3* | a | b | b | a | a | a |
|  | *hos15-2prr7-3* | a | b | b | a | ab | a |

*ANOVA and Tukey HSD test; *n* = 3

*Red highlights statistically significant differences

**Table S2. List of primers used in this study.**

| **Primer name** | **Sequence (5' -> 3')** | **Purpose** |
| --- | --- | --- |
| hos15-2 LP | TTCGAATATCCCTCCATTTCC | Mutant genotyping |
| hos15-2 RP | GCTGTTGTTTGGGACGTAAAG |  |
| PAC161-8474 | ATAATAACGCTGCGGACATCTACATTTT |  |
| prr7-3 LP | AGCAAGGACATACACTTTGGC |  |
| prr7-3 RP | TGAGAATTCGTCGTTCTTCAAC |  |
| LBb1.3 | ATTTTGCCGATTTCGGAAC |  |
| PRR9-1 LP | TTTCGTGGTTGTGATCGAAAG |  |
| PRR9-1 RP | AGGATCATCACGCAACTCATC |  |
| HA-HOS15 F (Kpn I) | ATGGTACCGATGTCTTCACTTACCTCCGT | Plasmid construction |
| HA-HOS15 R (Xho I) | ATCTCGAGTGCTACATTCTGAAATCAAGAACGC |  |
| GFP-PRR7 F (Sal I) | ACGTCGACTAATGAATGCTAATGAGGAGGG |  |
| TAP-PRR7 F (Sal I) | ATGCAGTCGACAATGAATGCTAATGAGGAGGG |  |
| TAP-PRR7 R (EcoR I) | GATCTGAATTCAGTTAGCTATCCTCAATGTTTTTTATGTC |  |
| HOS15_NT F (D-TOPO) | CACCATGTCTTCACTTACCTCC |  |
| HOS15_NT R noSTOP (D-TOPO) | AGATGTGGGAGTCATAACA |  |
| HOS15_CT F (D-TOPO) | CACCATGCAGACAAGTCACATTCCTAATTCTG |  |
| HOS15_CT R noSTOP (D-TOPO) | CATTCTGAAATCAAGAACG |  |
| CBF1 F (qRT-PCR) | GCATGTCTCAACTTCGCTGA | RT-qPCR |
| CBF1 R (qRT-PCR) | ATCGTCTCCTCCATGTCCAG |  |
| CBF2 F (qRT-PCR) | TGACGTGTCCTTATGGAGCTA |  |
| CBF2 R (qRT-PCR) | CTGCACTCAAAAACATTTGCA |  |
| CBF3 F (qRT-PCR) | GATGACGACGTATCGTTATGGA |  |
| CBF3 R (qRT-PCR) | TACACTCGTTTCTCAGTTTTACAAAC |  |
| COR15A F (qRT-PCR) | GGCCACAAAGAAAGCTTCAG |  |
| COR15A R (qRT-PCR) | CTTGTTTGCGGCTTCTTTTC |  |
| ACTIN2 F (qRT-PCR) | TAACAGGGAGAAGATGACTCAGATCA |  |
| ACTIN2 R (qRT-PCR) | AAGATCAAGACGAAGGATAGCATGAG |  |
| CBF-A_Fwd | TGCTTTCAAGGCCGAATGAT | ChIP-qPCR |
| CBF-A_Rev | CGTTCTCATTCCACGTGTGATG |  |
| CBF-B_Fwd | TTACCACTCTTTTTTTCCCTCTTTG |  |
| CBF-B_Rev | CTCGCTCTCACGTTATTGACATTT |  |
| CBF-C_Fwd | TCTTTACAAGGGTCAAAGGACACA |  |
| CBF-C_Rev | GCGAAGCAATCCCACGAT |  |
| CBF-D_Fwd | CGCGGAAACCATTGTCCATAC |  |
| CBF-D_Rev | GTACGAGACTCCACAGCGAG |  |
| CBF-E_Fwd | TGACTAAGGACGTGGTGGTTGA |  |
| CBF-E_Rev | AGCGCACTTCCTTCTCACTCA |  |
| CBF-F_Fwd | TGTTACATTTGATCATTCACCCAAA |  |
| CBF-F_Rev | CGTATATAAGCACGTAAGTCACCAAGT |  |
| CBF-G_Fwd | CGTGGCATTACCAGAGACACA |  |
| CBF-G_Rev | GCGGAAGATATTTTAGAGGCAAAA |  |
| CBF-H_Fwd | CAAGAGAGCACTGTCCGTAGCTT |  |
| CBF-H_Rev | TGGTTACAAGAGGAGCCACGTA |  |
| CBF-I_Fwd | GAGAGATGCTGGAAATTGTGATCA |  |
| CBF-I_Rev | AAATATGGTAAGTGGTTAGGCGAAA |  |
| CBF-J_Fwd | GGGTCAAAGGACACATGTCAG |  |
| CBF-J_Rev | GAACGCGGAGTTTCTGTCTC |  |
| COR15A-amp3_Fwd | ACAGAAACTCGGTAATGTGCA |  |
| COR15A-amp3_Rev | TGTTTGGAAATGAAAGGAGGAGT |  |
